# The alarmin Interleukin-33 modulates platelet proteome, function and biogenesis

**DOI:** 10.64898/2025.12.19.695476

**Authors:** Lucie Gelon, Stéphane Roga, Anne Gonzalez-de-Peredo, Edith Vidal, Jean-Philippe Girard, Sonia Severin, Emma Lefrançais

**Author notes:** Corresponding author : Emma Lefrançais, PhD IPBS CNRS 205 route de Narbonne 31044 Toulouse France **Text word count** 4000/4000.

## Abstract

Platelets, traditionally recognized for their involvement in hemostasis and wound healing, also play a central role in immune regulation and inflammation. Their function and production adapt in response to inflammatory cues such as cytokines and danger-associated molecular patterns. Interleukin-33 (IL-33), an alarmin released during tissue damage, particularly in lung inflammation, has been implicated in influencing platelet biology, though its exact effects remain poorly understood. To clarify IL-33’s role, we examined its impact on platelet production, proteome, adhesion, secretion, and aggregation using platelets from IL-33-deficient (IL-33KO) mice and IL-33 stimulation *in vivo*. Our results reveal that while platelets themselves do not express IL-33, platelets isolated from IL-33KO mice display altered proteomic signatures and reduced adhesion to fibrinogen, podoplanin, and laminin, alongside impaired thrombus formation under shear stress. IL-33 administration *in vivo* led to proteomic remodeling characterized by increased expression of inflammatory proteins, as well as changes in platelet morphology, including increased size, typically associated with de novo production. Using lung intravital microscopy, we visualized platelet fragmentation within the lung vasculature in real time, and observed enhanced fragmentation following IL-33 stimulation. Interestingly, ST2, the receptor for IL-33, is expressed in subsets of mouse and human megakaryocytes and hematopoietic progenitors, particularly those involved in a non-canonical pathway of thrombopoiesis that enables the rapid replenishment of platelets during inflammation, infection, and aging. Together, these findings identify IL-33 as a pivotal regulator of platelet function and production, linking inflammatory signaling to the dynamic regulation of thrombopoiesis.

**Key Points:** IL-33 impacts platelet inflammation-related proteins and adhesion

IL-33 receptor is expressed by subsets of MK progenitors in mouse and human

## Introduction

Platelets, small anucleated cell fragments derived from megakaryocytes (MKs), are essential for hemostasis and thrombosis. Beyond these roles, they have emerged as key players in immune responses and inflammation, particularly in lung diseases such as asthma, ARDS, and pulmonary fibrosis, through interactions with leukocytes and endothelial cells, and by releasing soluble mediators^1–3^. Interleukin-33 (IL-33), a nuclear alarmin abundantly expressed in epithelial cells of barrier tissues, endothelial cells, and fibroblastic stromal cells in various tissues, is a potent inducer of type 2 immune responses^4–6^. It plays a central role in initiating inflammation and tissue repair, particularly in asthma and other pulmonary disorders. IL-33 activates resident immune cells including ILC2s, Tregs, and mast cells, contributing to airway inflammation and remodelling^7^. It also affects hematopoiesis, such as hematopoietic stem cell mobilization^8,9^, and has been associated with inhibition of erythropoiesis^10^. Recent studies suggest IL-33 may regulate platelet biology. IL-33 overexpression has been linked to thrombocytosis^11^ and in models of intestinal^12^ and allergic lung inflammation^13,14^, IL-33-deficient mice showed altered platelet activation and reduced neutrophil and eosinophil recruitment, pointing to a potential role of platelet-derived IL-33 in pulmonary inflammation. However, IL-33 expression by platelets remains controversial. IL-33 is not typically expressed in hematopoietic cells^5,6^ and has not been consistently detected by transcriptomic or proteomic analyses under homeostatic or inflammatory conditions^15,16^. Single-cell analyses have revealed transcriptional heterogeneity in MK populations, identifying distinct subsets with specialized functions^17,18^. A low-ploidy MK population, potentially involved in immune regulation, has been described in lung, bone marrow, liver, blood, and spleen^19–22^. Notably, the lung environment, and specifically IL-33, has been implicated in the maturation of pulmonary MKs, potentially influencing their immune phenotype^23^. Together, these findings suggest that IL-33 could influence platelet production, content, and function via effects on subsets of MKs. To explore this hypothesis, we used IL-33 knockout mice and *in vivo* stimulation with recombinant IL-33 to assess its impact on platelet biogenesis, proteome, and functional properties.

## Methods

### Platelet and megakaryocyte Immunofluorescence

200 µL of platelet suspension (3×10⁶) or 100 µL of megakaryocyte-enriched solution was prepared as detailed in Supplemental methods and deposited on poly-L Lysine coated coverslips. After 30 min sedimentation (RT), cells were fixed (4% PFA, 10 min), permeabilized (0.5% Triton, 10 min), and blocked (3% BSA, 30 min). Primary antibodies (in 0.3% BSA) were incubated overnight at 4 °C: anti-IL-33 (1:200), phalloidin-CF488A (1:400), APC-CD41 (1:1000). After PBS washes, Cy3 anti-goat secondary antibody was added for 1 h (RT). Slides were mounted with Mowiol/DAPI.

### Western Blot analysis

Samples were lysed in RIPA buffer (with protease inhibitors) under agitation at 4 °C for 30 min–2 h, then centrifuged (12,000 rpm, 20 min, 4 °C). Supernatants were mixed 1:1 with 2X Laemmli buffer and boiled (95 °C, 5 min). 20 µg of protein was loaded in each lane and run on 12% SDS-PAGE. Proteins were transferred to nitrocellulose (90 V, 1 h, 4 °C), blocked in 5% milk/TBS-T (1 h, RT), and incubated overnight at 4 °C with anti-IL-33 (R&D AF3626) in 2.5% milk/TBS-T. After TBS-T washes, membranes were incubated (1 h, RT) with HRP-conjugated anti-goat IgG (Promega V8051), washed, and developed using ECL Prime (GE) and ODYSSEY XF (Licor).

### Platelet activation assays and fibrinogen binding assays

Platelet-rich plasma (PRP), diluted to 3000 platelets/µL, was stimulated for 15 min at room temperature in the dark with ADP (10 µM), CRP (5 µg/mL), or thrombin (0.01 U/mL). Samples were stained for 15 min at room temperature in the dark (see Supplemental Table 1), centrifuged (600g, 7 min, 20 °C), resuspended in Tyrode’s buffer, and analyzed on a Fortessa X20 flow cytometer. For soluble fibrinogen binding, PRP was activated or not with ADP (1 µM, 15 min, RT, dark) and incubated with AF488-fibrinogen (20 µg/mL, 15 min, RT, dark), centrifuged (600g, 7 min, 20 °C), resuspended in Tyrode’s buffer, and analyzed by Fortessa X20 cytometry.

### Platelet binding and spreading assay

Lab-Tek slides were coated overnight at 4 °C with fibrinogen (200 µg/mL), podoplanin (2 µg/mL), laminin (50 µg/mL), or vitronectin (10 µg/mL), washed and blocked with 1% BSA (1–2 h, RT). 12,000 platelets/µL were unstimulated or stimulated with ADP (1 µM, 10 min), centrifuged (600g, 7 min, 20 °C), and resuspended in Tyrode’s buffer. Platelets were seeded on coated slides for 5 or 15 min at 37 °C ± apyrase (2 U/mL), washed, fixed (4% PFA, 10 min), blocked (3% BSA, 30 min, RT), stained with CD41-APC (1:400 in 0.3% BSA, 1 h, RT), washed, mounted with Mowiol, and imaged on a ZEISS Imager.M2 AXIO (×40 objective).

### Thrombus formation under flow assay

Vena8 Fluoro+ biochip microcapillaries were coated with collagen (50 µg/mL, 1 h) and blocked with 0.5% fatty acid-free BSA. Intracardiac blood was collected with a heparinized syringe (100 µL, 100 U/mL), labeled with DIOC6 (2 µM, 5 min, 37 °C), and perfused through collagen-coated channels at shear rates of 1500 s⁻¹ (homeostatic) and 3000 s⁻¹ (pathological) using a syringe pump. Thrombus formation was recorded in real time via 40x oil immersion microscopy and analyzed with Imaris 9.3.

### Hematoanalysis

50 µL of blood were collected from the mandible into an ACD-coated Minivette, and 30 µL was analyzed with an automated hematology analyzer (PE-7010vet 5 Diff, PROKAN).

### Platelet release in lung capillaries observed by lung intravital microscopy

Lung intravital microscopy was performed in PF4-mTmG mice 24 h after one or three daily intranasal doses of rIL-33 (1 µg). Using the intercostal thoracic window method^24,25^, mice were anesthetized, ventilated (tidal volume 10 µL/g, 130–140 breaths/min, PEEP 2–3 cm H₂O), and hydrated hourly. The left lung lobe was stabilized by suction (20–25 mmHg) and imaged with a Zeiss 7 MP two-photon microscope (20×/1.0, 920 nm) every minute for 60–120 min. Images were analyzed in Imaris to quantify MKs and proplatelet release, classified by size: Small (<500 platelets), Medium (500–1000), Large (>1000), as previously described^21^.

## Results

### IL-33 is not expressed by circulating platelets

IL-33 protein expression in platelets and megakaryocytes has been reported^13^. However, IL-33 mRNA and protein have not been detected in previous platelet transcriptomic or proteomic studies^26,27^. To resolve these discrepancies, we performed analyses with a polyclonal goat anti-mouse IL-33 antibody (R&D AF3626) that has been validated for immunofluorescence, flow cytometry and western blot analyses in previous studies (**Supplemental Table 1**), including several studies by our team^6,28–30^. IL-33 is a nuclear cytokine and the Ab AF3626 stains the nucleus of IL-33 producing cells in WT mice but not in IL-33KO mice. These include epithelial cells, endothelial cells and stromal cells in various tissues, and astrocytes/oligodendrocytes in the brain (see references in **Supplemental Table 1**). Although AF3626 antibody specifically detects endogenous nuclear IL-33 protein in *bona fide* IL-33-producing cells, it can also give non-specific cytoplasmic staining in some cells, for instance, cells of the lamina propria in the colon^6^. In the current study, we observed cytoplasmic staining of bone marrow-derived megakaryocytes with AF3626 antibody (**Fig. 1A**) but the staining was non-specific (still present in IL-33KO megakaryocytes). Similarly, we observed staining of WT platelets with AF3626 antibody by both immunofluorescence (**Fig. 1B**) and flow cytometry (**Fig. 1C**), but the staining was the same with IL-33KO platelets (**Fig. 1B, 1C**). Thus, we could not detect expression of endogenous IL-33 protein in WT megakaryocytes and platelets by immunofluorescence and flow cytometry techniques. Western blot analyses using total lung lysates as positive controls confirmed that endogenous IL-33 protein is not expressed by WT megakaryocytes and platelets (**Fig. 1D**). Finally, flow cytometry analysis of cells from IL-33-Citrine reporter mice revealed that the IL-33 promoter is active in CD41^-^CD45^-^ bone marrow stromal cells (**Fig. 1E, 1F**)^31^ but not in CD41^+^ megakaryocytes and platelets (**Fig. 1E**). Together, these experiments provide strong evidence that endogenous IL-33 protein is not expressed in megakaryocytes and platelets, and that previously reported signals likely reflect antibody cross-reactivity rather than IL-33 expression.

**Figure 1.**
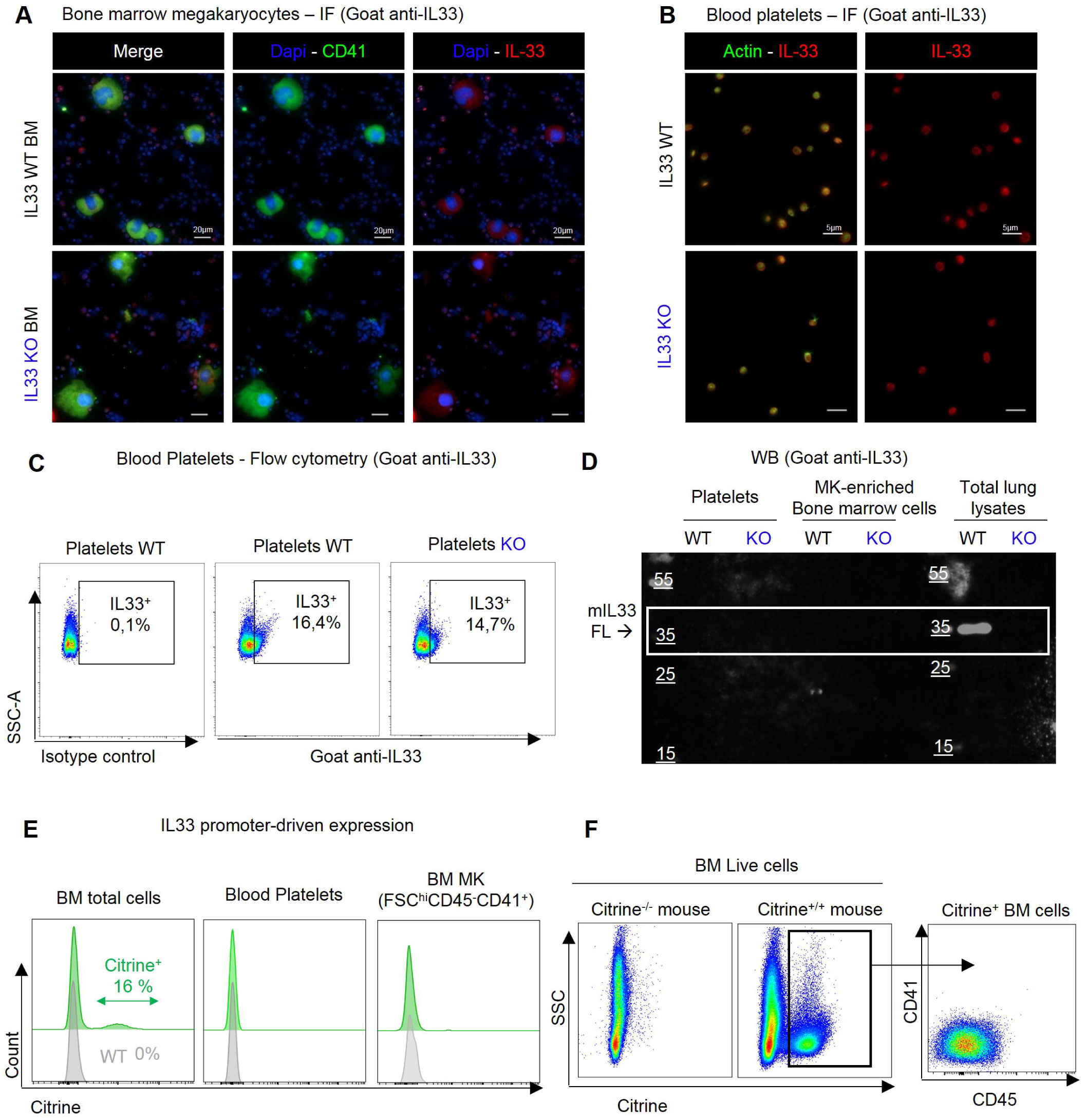
IL-33 is not expressed by circulating platelets or megakaryocytes. (A–B) Representative immunofluorescence images showing nonspecific IL-33 staining in bone marrow–derived megakaryocytes enriched on BSA gradient (A) and platelet-rich plasma (PRP) (B) from WT and IL-33KO mice. (C) Flow-cytometric detection of IL-33 nonspecific signal in blood platelets from WT and IL-33KO mice (PRP), using an isotype control as a negative reference. (D) Western blot analysis of IL-33 expression in platelets and megakaryocyte-enriched bone marrow cells, with total lung lysates from WT and IL-33 KO mice used as positive and negative controls, respectively. Following quantification, an equal amount of protein (20 µg) was loaded in each lane (E) IL-33 promoter activity in total bone marrow cells, blood platelets and BM MK from IL-33-Citrine reporter mice assessed by flow cytometry. (F) Identification of Citrine⁺ cells as stromal cells (CD41⁻CD45⁻) in Citrine reporter mice (Citrine^+/+^) by flow cytometry. Panels A–D used a polyclonal goat anti-mouse IL-33 antibody (R&D Systems, AF3626).

#### IL-33 receptor is expressed by subsets of megakaryocytes and hematopoietic progenitors

To assess whether IL-33 acts directly on platelets or their progenitors, we analyzed ST2 expression via flow cytometry and found it absent in platelets (**Supplemental Fig. 1A**). ST2, however, is known to be expressed in hematopoietic progenitors and has been implicated in HSC mobilization and erythroid inhibition^8–10^. Recent studies have identified resident and immature pulmonary megakaryocytes in mice, which differ from bone marrow megakaryocytes by their expression of genes related to innate immunity and inflammation^21–23^. Among the 543 genes overexpressed in these pulmonary megakaryocytes, we identified the IL-33 receptor (*Il1rl1/St2*)^21^ (**Fig 2A**) and confirmed its protein expression by flow cytometry (**Fig 2B**). Lung megakaryocytes arise from non-canonical fast-track megakaryopoiesis^32^, differentiating from CD41^+^ MK-biased HSC^33,34^ and non canonical CD48^-/low^ megakaryocyte progenitors (MkP)^35–37^, bypassing multipotent progenitors. This pathway ensures platelet supply under stress (acute thrombocytopenia, inflammation^38^, lung infection^39^) and is increased during myelofibrosis^40^ and ageing^37^. Given that non-canonical megakaryopoiesis is enhanced with age, we analyzed LT-HSCs and MkPs from young and aged mice by flow cytometry (**Supplemental Fig. 2A-B**) and measured ST2 expression on these populations (**Fig. 2C** and **Supplemental Fig. 2C**). Mk-biased LT-HSCs (**Fig. 2D-E**) and MkPs (**Supplemental Fig. 2D-E**) expanded with age, consistent with previous studies ^37,41^. Notably, ST2 expression rose significantly in aged Mk-biased LT-HSCs, with ∼20% ST2⁺ cells (**Fig. 2F-G**), and was also elevated in MkPs (**Supplemental Fig. 2F-G**). In BM MKs from young and aged mice, ST2 expression was heterogeneous. We examined correlations between ST2 mean fluorescence intensity (MFI) and MK markers, finding negative correlations with cell granularity (SSC), CD9, CD41, and CD45 (**Supplemental Fig. 2H**). These results suggest that elevated ST2 expression marks a less mature, progenitor-like MK phenotype. Finally, human single-cell RNA-seq data indicate ST2 enrichment in hematopoietic progenitors biased toward erythroid, eosinophil, mast cell, and megakaryocyte lineages in bone marrow, spleen, and blood^42^ (**Fig. 2H**). Collectively, these findings indicate that ST2 is expressed by medullary and extramedullary specific subsets of immature megakaryocytes and hematopoietic progenitors, suggesting that IL-33 may influence platelet biology, particularly under conditions that engage non-canonical megakaryopoiesis.

**Figure 2.**
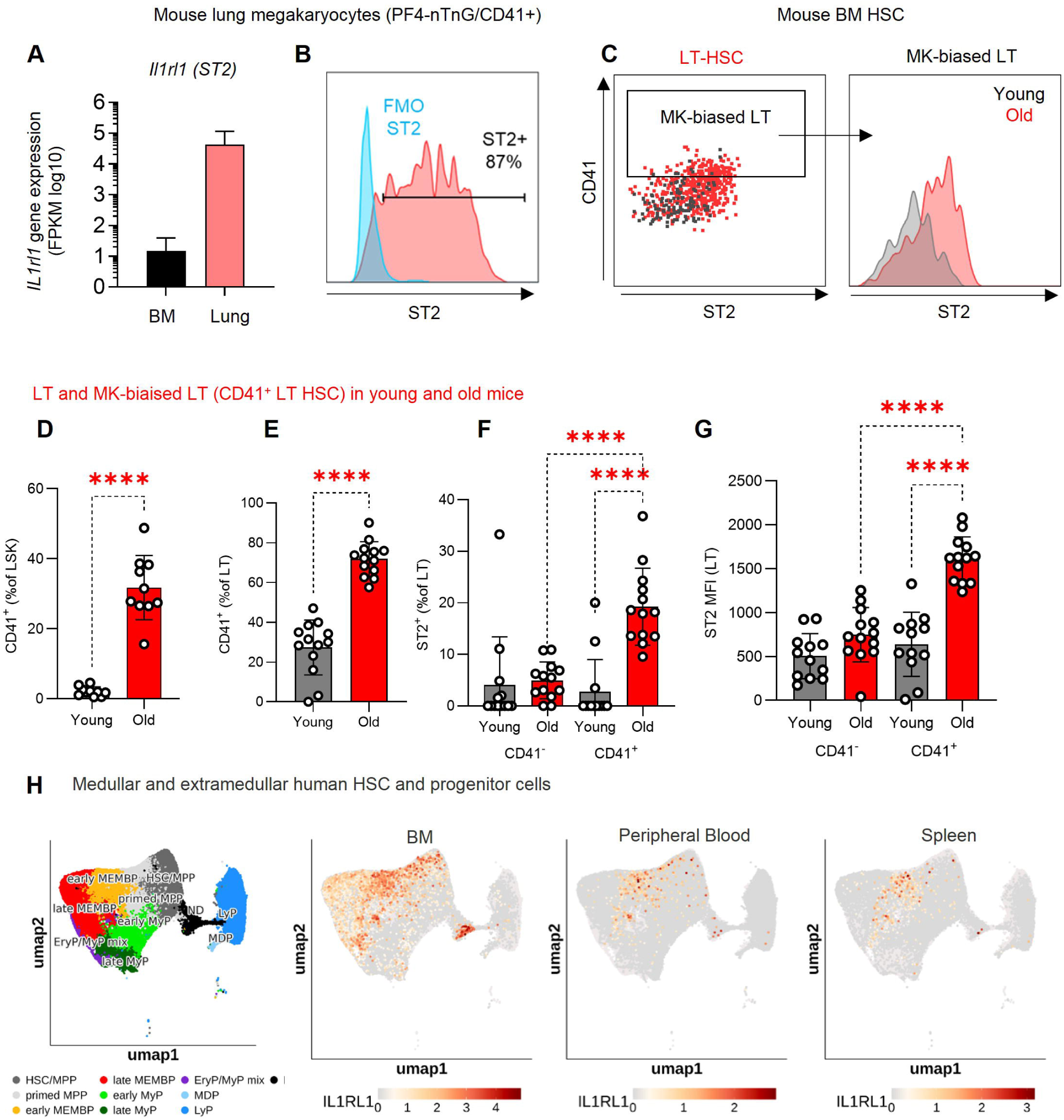
ST2 expression in subsets of hematopoietic and megakaryocyte progenitors associated with non-canonical hematopoiesis in mouse and human (A) Comparative *st2* (*il1rl1*) gene expression analysis (RNA-seq) of murine bone marrow versus lung megakaryocytes (PF4-nTnG/CD41⁺ cells). Data from Lefrançais et al., Nature, 2017 (n=3 mice per group). (B) Detection of ST2 (IL1RL1) protein expression in lung megakaryocytes (PF4-nTnG/CD41⁺ cells) by FACS. (C) Gating strategy for identifying megakaryocyte-biased LT-HSCs (CD41⁺ LT-HSCs) and analysis of ST2 expression in young (8-week-old) and aged (2-year-old) mice. (D–E) Quantification of MK-biased LT-HSCs in young and aged mice. (F–G) Quantification of ST2⁺ cells and ST2 expression levels within LT-HSCs (F–G). (D-G) Data obtained from n = 13 mice per group (biological replicates) across three independent experiments conducted on separate collection and analysis days. Investigators were blinded to mouse age during bone marrow collection and analysis. (H) IL1RL1 expression in single-cell analysis of human HSPCs from bone marrow, peripheral blood, and spleen. (source: https://bioinf.stemcells.cam.ac.uk/shiny/laurenti/ExtramedHSPCs/).

### IL-33 deficiency in vivo does not affect platelet count or morphology

Since IL-33 receptors are expressed on subsets of megakaryocytes and hematopoietic progenitors, we investigated whether IL-33 deficiency affects platelet count under steady-state conditions. Platelet number and size were analyzed in WT and IL-33KO mice using flow cytometry and hematological analysis. No significant differences were observed in platelet count, size, or granularity between the two groups (**Supplemental Fig. 3A–E**). These findings suggest that IL-33 is not required for maintaining platelet count or baseline platelet morphology under homeostatic conditions.

### IL-33 deficiency alters the platelet proteome

To investigate the impact of IL-33 on platelet biology and better understand the functional defects observed with platelets from IL-33 KO mice^12,13^, we performed a proteomic analysis of isolated platelets from WT and IL-33KO littermates using mass spectrometry (**Fig. 3A**). A total of 1583 proteins were detected, with 129 upregulated and 36 downregulated in platelets from IL-33KO mice (**Fig. 3B-C, Supplemental Tables 2, 5 and 6**). Gene ontology analysis revealed enrichment in processes related to actin cytoskeletal remodelling, and lamellipodium assembly (**Fig. 3D-E**), with increased expression of key cytoskeletal regulators such as CDC42, Pleckstrin, Coronin, Rac1, and RhoA (**Supplemental Table 2)**. In contrast, several pathways involved in platelet activation and adhesion were significantly downregulated (**Fig. 3F-G**).

**Figure 3.**
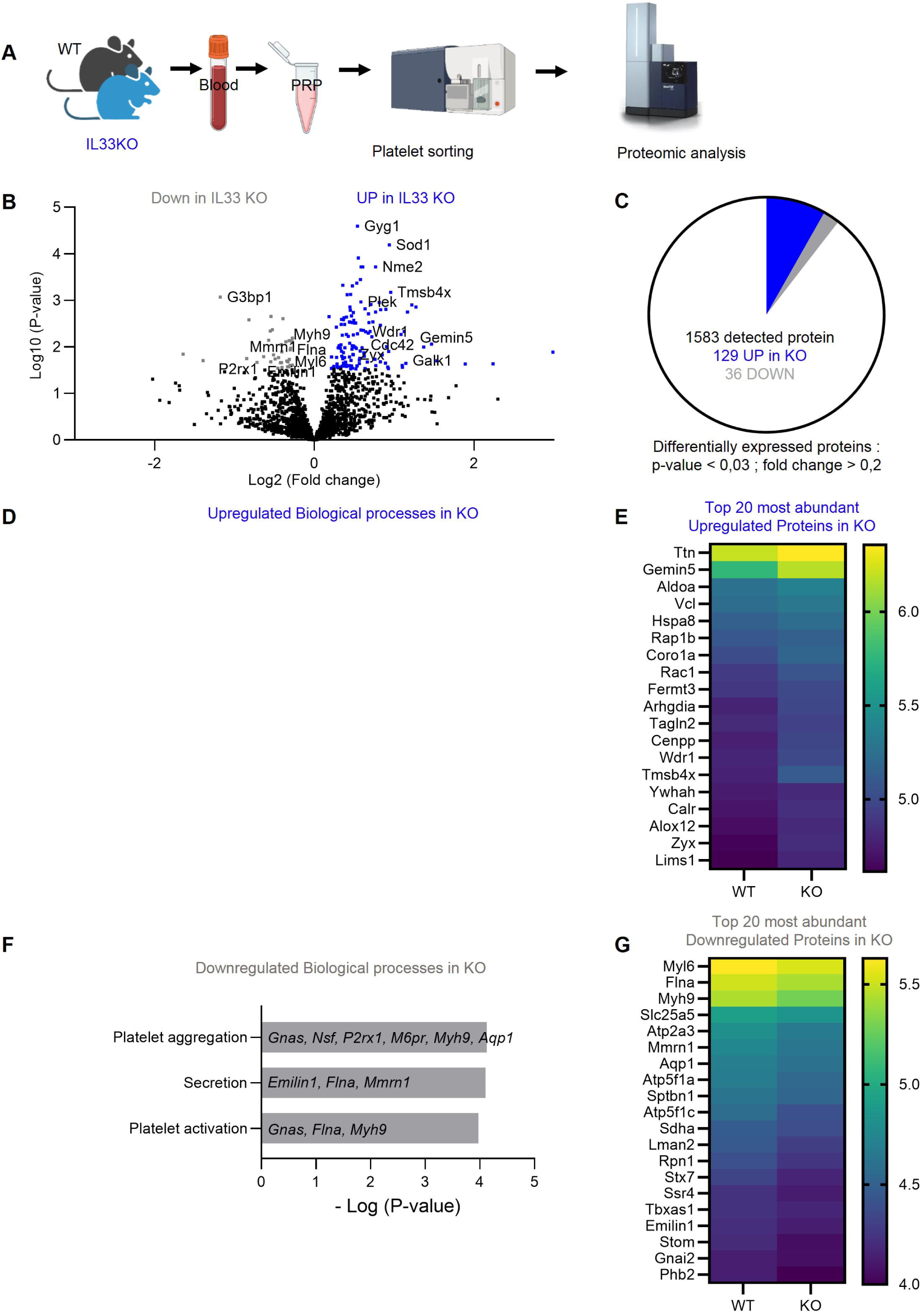
IL-33 deficiency alters the platelet proteome (A) Proteomic profiling of sorted platelets from WT and IL-33KO mice. (B) Volcano plot comparing IL-33KO and WT platelet proteomes. The x-axis shows log fold change (IL-33KO/WT), and the y-axis shows –log(p-value). Proteins upregulated in IL-33KO platelets are highlighted in blue; downregulated proteins in grey. (C) Number of proteins detected and significantly altered (p < 0.03, fold change > 0.2). (D, F) Gene Ontology (GO) analysis of enriched biological processes among upregulated (D) and downregulated (F) proteins (https://geneontology.org). (E, G) Heatmaps of the top 20 most abondant upregulated (E) and downregulated (G) proteins. Data were obtained from n = 5 mice per group.

Notably, important extracellular matrix-binding proteins and cytoskeletal components, including Emilin1, Multimerin-1 (MMRN1), Filamin A, and Myosin heavy chain 9 (Myh9), were downregulated in platelets from IL-33KO mice (**Supplemental Table 2)**. Flow cytometry analysis further showed no major differences in the expression of most integrins and receptors between WT and KO platelets, except for decreased CD49f, involved in laminin binding and increased CD150 in platelets from IL-33KO mice (**Supplemental Fig. 4A-B**).

### IL-33 deficiency impairs platelets adhesion to soluble fibrinogen and extracellular matrix components

Given the proteomic alterations in cytoskeletal and adhesion-related proteins, we investigated the adhesive properties of platelets from IL-33KO mice. First, we evaluated their ability to bind soluble fibrinogen following ADP activation (**Fig. 4A**) and observed a reduced binding capacity in platelets from IL-33KO mice (**Fig. 4B**). Under static conditions, we evaluated platelet adhesion and spreading on fibrinogen, podoplanin, and laminin-coated surfaces (**Fig. 4C**). After ADP stimulation, platelets from IL-33KO showed reduced adhesion to all substrates (**Fig. 4D-F)** along with impaired spreading on fibrinogen, evidenced by a higher proportion of rounded platelets (**Fig. 4D**). To determine whether this defect was ADP-dependent, we repeated adhesion assays without ADP (**Fig. 4G-H**) or with ADP blocked by apyrase (**Supplemental Fig. 5A**). Platelets from IL-33KO mice displayed reduced binding to fibrinogen and podoplanin at 5 and 15 minutes under both conditions (**Fig. 4G-H, Supplemental Fig. 5B-C**). These findings indicate that IL-33 deficiency impairs platelet adhesion and spreading, particularly on extracellular matrix components.

**Figure 4.**
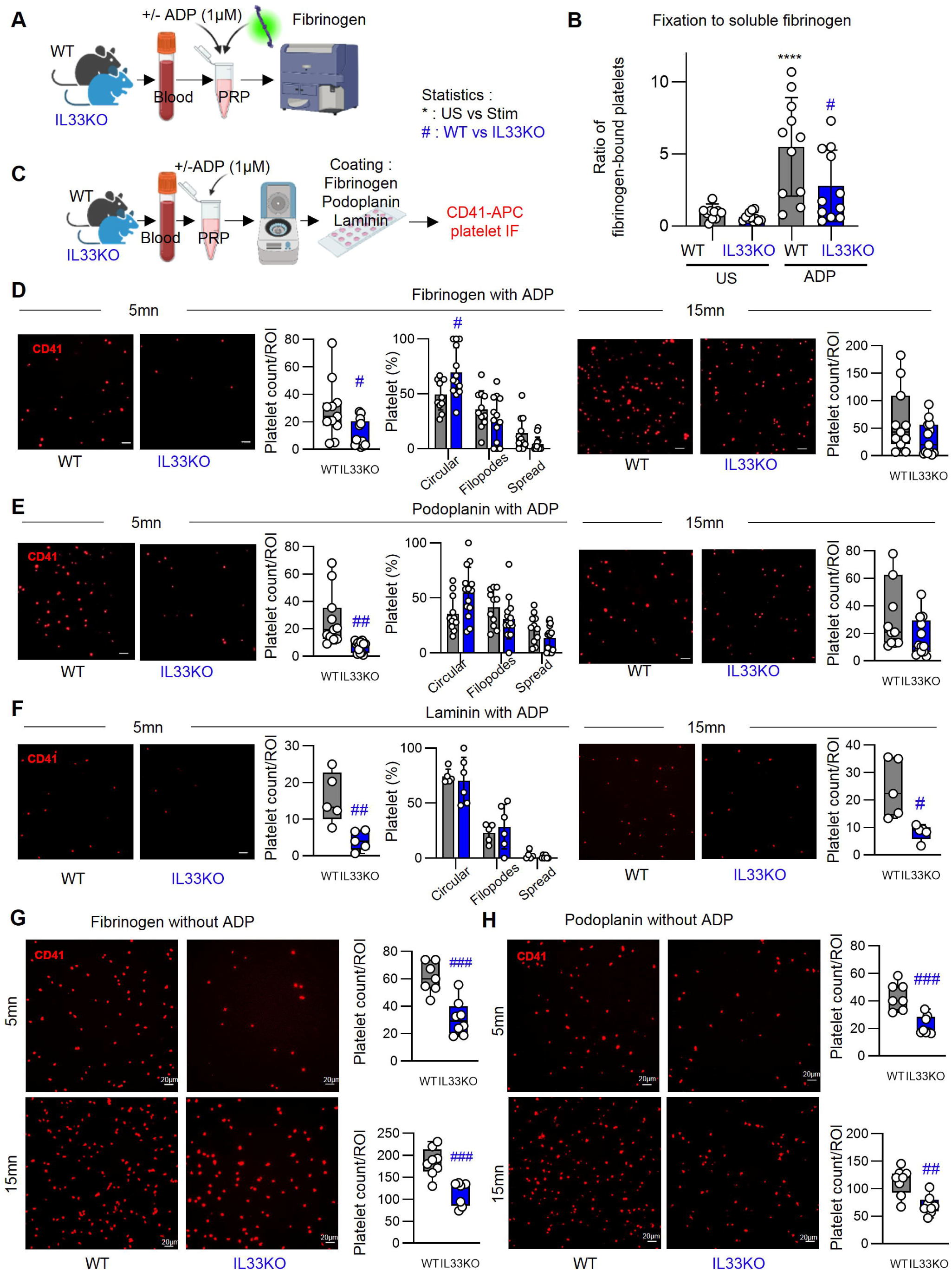
IL-33 deficiency impairs platelets adhesion to extracellular matrix components (A) Schematic and (B) analysis of platelet adhesion assays to soluble fibrinogen, with or without ADP (1 µM), assessed by flow cytometry. Platelet-rich plasma (PRP) from WT and IL-33 KO mice was analyzed, and results are expressed as a ratio relative to the WT mean. Data were obtained from n = 11 mice per group across 3 independent experiments. (C) Schematic of platelet spreading assays on fixed substrates, analyzed by immunofluorescence in WT and IL-33 KO platelets. Representative images and quantitative analyses of adherent and spreading PRP from WT or IL-33 KO mice on (D) fibrinogen, (E) podoplanin, and (F) laminin at 5 and 15 minutes following ADP (1 µM) stimulation. Platelets were stained with CD41-APC. Data were obtained from n = 11 mice per group across three independent experiments, except for the laminin spreading assay, which was performed on n = 5 mice in a single experiment. (G-H) Platelet spreading assays without ADP stimulation on fibrinogen (G) and podoplanin (H) at 5 and 15 minutes (n = 7 mice per group per experiment). Statistical analyses were performed using t-tests (D, E, F) and two-way ANOVA with Sidak’s multiple comparisons test (B). Scale bar = 20 µm. t-test, #p < 0.05; ##p < 0.01; ###p < 0.001; ####p < 0.0001.

### IL-33 deficiency reduces thrombus formation on collagen under high shear stress

To assess the impact of IL-33 deficiency on thrombus formation, we conducted *ex vivo* thrombus formation assay. Whole blood from WT or IL-33 KO mice was perfused through collagen-coated microcapillaries under physiological or pathological wall shear rates of 1500 and 3000 s^−1^, respectively (**Fig. 5A**). Under physiological arterial shear rate (1500 s^-1^; 60 μL/min), thrombus formation was similar between platelets from WT and IL-33KO mice (**Fig. 5B**). However, under pathological arterial flow conditions (3000 s^-1^; 120 μL/min), platelets from IL-33KO mice formed significantly smaller thrombi compared to WT platelets (**Fig. 5C**). These findings suggest that IL-33 contributes to proper platelet adhesion to collagen and supports platelet function under high shear stress.

**Figure 5.**
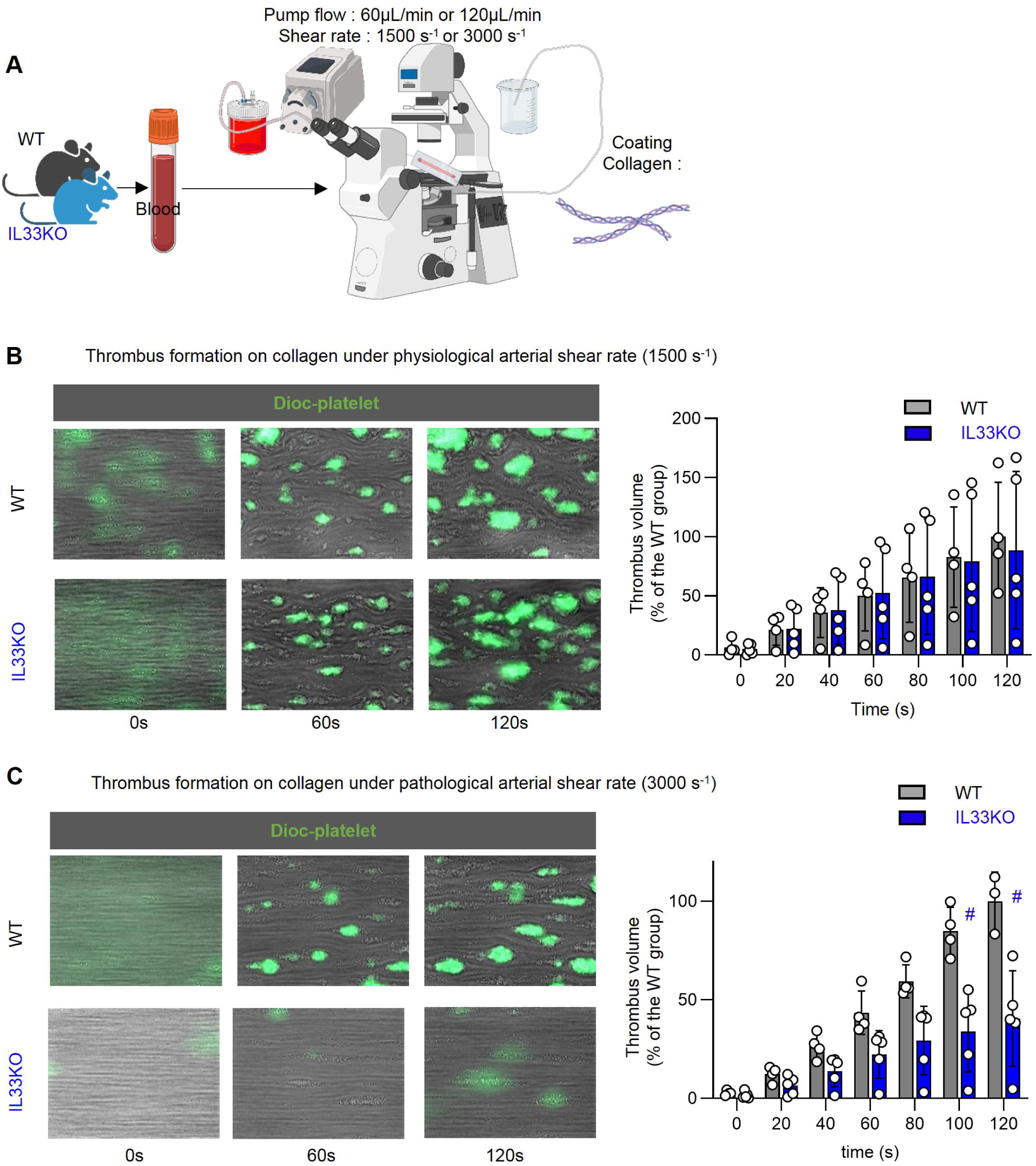
IL-33 deficiency reduces thrombus formation under high shear stress (A) Schematic representation of the thrombus formation assay on collagen during arterial flow of whole blood from WT or IL-33 KO mice. (B) Representative images and quantitative analysis of thrombus volume, expressed as a percentage relative to the WT mean, formed on collagen under physiological arterial shear rate (1500 s⁻¹) using whole blood from WT or IL-33 KO mice. (C) Representative images and quantitative analysis of thrombus volume, expressed as a percentage relative to the WT mean, formed on collagen under pathological arterial shear rate (3000 s⁻¹) using whole blood from WT or IL-33 KO mice. Data are representative of three independent experiments (n = 4-5 mice/group). Statistical analysis was performed using two-way ANOVA with Sidak’s multiple comparisons test, #p < 0.05. Investigators were blinded to genotype during the thrombus formation assay.

### IL-33 deficiency affects αIIbβ3 integrin activation and CD40L expression upon stimulation without impairing degranulation or aggregation

To evaluate the impact of IL-33 deficiency on platelet activation and degranulation, we stimulated platelets from IL-33KO mice with the soluble agonists ADP, CRP, and thrombin (**Supplemental Fig. 6A**). P-selectin and CD63 surface expression were unchanged (**Supplemental Fig. 6B-E**) indicating intact alpha and dense granule release. However, thrombin-induced CD40L expression was reduced in platelets from IL-33KO mice (**Supplemental Fig. 6D**), as was αIIbβ3 integrin activation measured by Jon/A binding (**Supplemental Fig. 6F-G**). Despite this, platelets from IL-33KO mice displayed normal aggregation in response to collagen, CRP, and thrombin (**Supplemental Fig. 7A-B**). These results suggest that IL-33 deficiency impairs αIIbβ3 integrin activation without broadly affecting degranulation or aggregation. The previously observed defects in binding capacity may result from insufficient activation of the αIIbβ3 integrin.

### IL-33 treatment in vivo does not alter platelet count but affects platelet morphology and activation state

IL-33 overexpression in mice can induce thrombocytosis^11^, and its receptor ST2 is expressed on megakaryocyte progenitors. Surprisingly, IL-33 deficiency alters the platelet proteome and reduces adhesion at baseline, despite IL-33 typically being released in response to tissue injury, revealing a physiological role for IL-33 expressed by stromal cells from birth. To model an acute response to inflammation or injury and assess effects on platelet count and function, wild-type mice were treated intranasally with recombinant IL-33 (**Fig. 6A**). Platelet counts were unchanged (**Fig. 6B**), but treated mice showed increased platelet size and granularity (**Fig. 6C**), together with a transient rise in Thiazole orange (TO)⁺ immature platelets from 1 h to 18 h after IL-33 administration, which returned to baseline by 24 h (**Supplemental Fig. 8A**). These observations are consistent with a short-lived pulse of de novo thrombopoiesis^43^. We also observed enhanced degranulation and activation, marked by higher P-selectin, CD63 (**Fig. 6D**) and plasma sCD40L (**Fig. 6E**). The increased platelet size observed after IL-33 treatment suggests that active thrombopoiesis is ongoing, potentially compensating for platelet activation and consumption to maintain platelet count.

**Figure 6.**
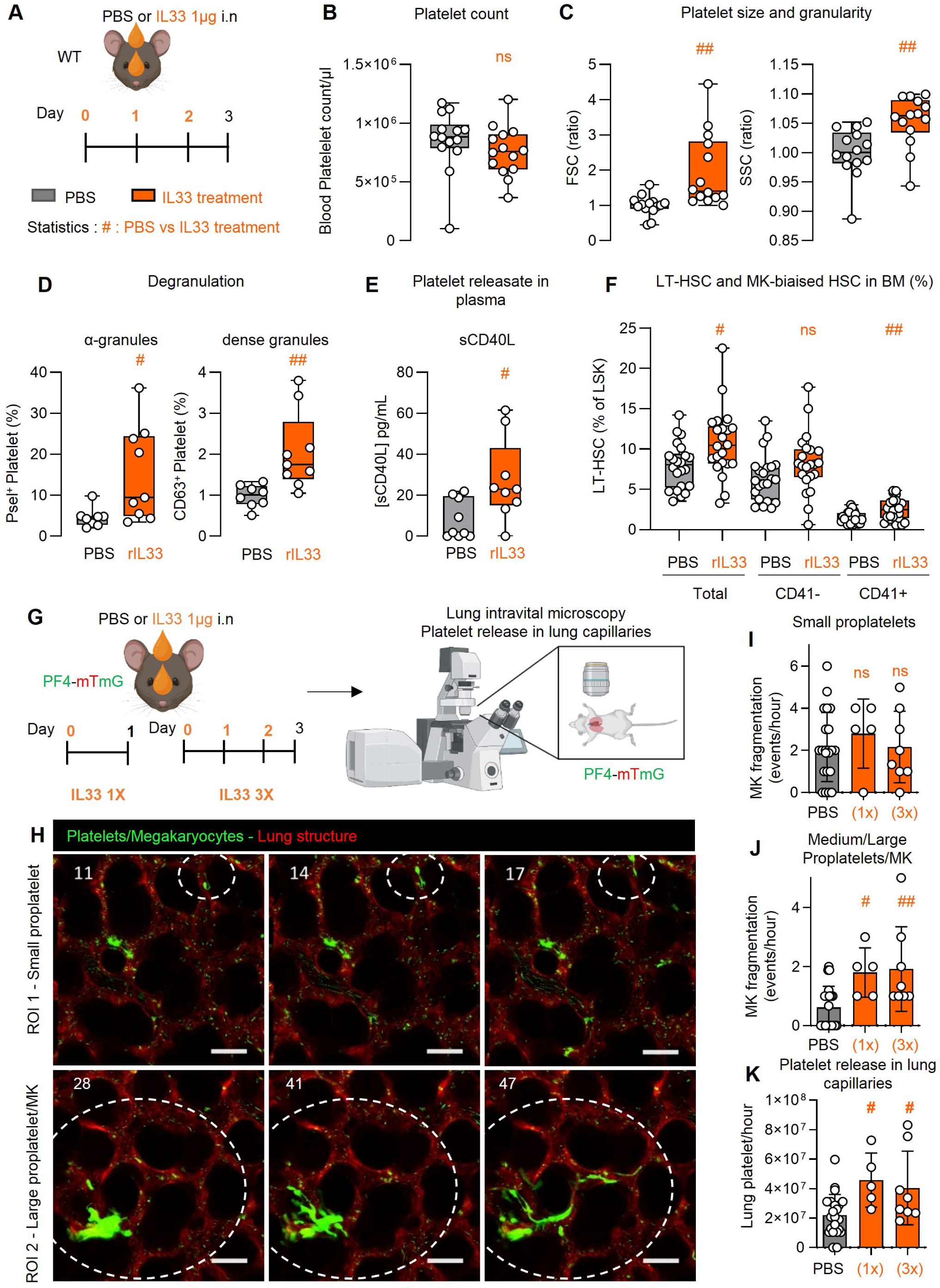
IL-33 treatment *in vivo* modifies platelet morphology and activation, expands the megakaryocyte-biased LT-HSC pool, and enhances platelet release within lung capillaries. (A) Schematic overview of the IL-33 treatment protocol: WT mice received three intranasal doses of recombinant human IL-33 or PBS. (B) Flow cytometry analysis of platelet morphology in WT mice following IL-33 or PBS treatment. (C) Quantification of platelet counts by flow cytometry in IL-33- or PBS-treated WT mice. Data are from n = 14 mice per group, collected across three independent experiments. (D) Flow cytometry assessment of platelet degranulation, measuring alpha granule (P-selectin) and dense granule (CD63) expression after IL-33 or PBS administration. (E) Plasma levels of sCD40L, measured by ELISA in IL-33- or PBS-treated WT mice. (D-E) Data are from n = 8-9 mice per group, collected across 2 independent experiments. (F) Quantification of bone marrow long-term hematopoietic stem cells (LT-HSC), separated into CD41⁻ (canonical) and CD41⁺ (megakaryocyte-biased) populations, shown as a percentage of LSK cells. Data are from *n* = 22 mice per group, collected across 5 independent experiments. (G) Lung intravital microscopy was performed in PF4-mTmG mice treated with IL-33 or PBS, 24 hours after the first (IL-33 1x) or third (IL-33 3x) administration to assess megakaryocytes/proplatelets generating platelets. (H) Representative time-lapse intravital lung microscopy images showing PF4⁺ small proplatelets and large proplatelet/megakaryocyte structures releasing platelets within pulmonary capillaries. Scale bar = 50 µm. (I–J) Quantification of proplatelet structures producing platelets per hour per ROI in the lung, categorized as small (<200 platelet volume equivalent), medium (200–1000 platelets), and large (>1000 platelets). (K) Total platelet release per hour across the entire lung. Statistical analysis: t-test, #p < 0.05, ##p < 0.01. A total of *n* = 5–21 videos were analyzed, collected from 3–11 mice per group across five independent experiments.

### IL-33 treatment in vivo increased MK biaised LT-HSC and platelet release in lung capillaries

Thrombopoiesis involves two key stages: megakaryocyte differentiation and platelet release. Given that ST2 is expressed on MK-biased hematopoietic stem cells (HSCs) and lung-resident MKs, both associated with non-canonical thrombopoiesis, we examined the impact of IL-33 on these populations, as well as on platelets generated in the lung vasculature from circulating MKs^21^, a process known to intensify under acute demand^44,45^. Following IL-33 administration, we observed an expansion of MK-biased LT-HSCs (**Fig. 6F**) whereas MkP populations remained unchanged (**Supplemental Fig. 8B-C**). In parallel, using lung intravital microscopy and platelet reporter mice, we detected increased numbers of medium to large proplatelets and MKs actively releasing platelets within the pulmonary vasculature (**Fig. 6G-K, Supplemental Fig. 8D**, **Supplemental Video 1**). We previously showed that circulating whole megakaryocytes (MKs) and large proplatelets detected in the lung originate from the bone marrow ^21^. Their egress and subsequent release in the lung vasculature is associated with rapid and more efficient platelet production^46^. In mice, large MKs typically fragment within less than one hour (∼60000 µm³ per hour) in lung capillaries (**Supplemental Video 1**)^21^, whereas in the bone marrow, fragmentation occurs at a slower rate (∼2000 µm³ per hour)^47^, requiring about 30 hours for a mature 50 µm MK to fully fragment. Together, these findings support a role for IL-33 in thrombopoiesis by expanding MK-biased HSCs in the bone marrow and accelerating platelet release in the lungs.

### In vivo IL-33 treatment does not impact the platelet’s ability to form thrombi but does alter the platelet proteome by modulating pathways related to inflammation and coagulation

To better understand if IL-33 stimulation *in vivo* also affects platelets function, we analysed both their proteome and thrombus formation capacity. Using blood from wild-type (WT) mice treated with PBS or recombinant IL-33, thrombus formation on collagen was assessed under shear rates of 1500 s⁻¹ and 3000 s⁻¹ (**Supplemental Fig. 9A**). Thrombus volume was comparable between the two groups under both conditions (**Supplemental Fig. 9B-C**), indicating that IL-33 stimulation *in vivo* does not affect thrombus formation under flow. The impact of acute IL-33 stimulation in adult animals has thus distinct effects compared to lifelong IL-33 deficiency. We then performed a global proteomic analysis of platelets from WT mice treated with PBS or recombinant IL-33. Out of 1,072 proteins identified, 58 were upregulated and 64 downregulated (**Fig. 7A-C, Supplemental Tables 7 and 8**). Gene ontology analysis revealed that rIL-33 treatment led to an enrichment of immune-related processes and cell adhesion pathways (**Fig. 7D**). IL-33 upregulated inflammatory and adhesion associated mediators (**Fig. 7E, Supplemental Tables 2 and 7**), including PPBP (CXCL7), which promotes granulocyte recruitment and tissue repair^48^; LGALS3 (galectin-3), which binds platelet GPVI and facilitates cell extravasation^49^ and HS1 (HCLS1), a GPVI-dependent adaptor involved in cytoskeletal reorganization^50^. Another notable hit is MMP12 (matrix metalloproteinase-12), as it is also increased in lung versus bone marrow megakaryocytes (**Supplemental Figure 10**) and links platelet activation, hematopoiesis, and lung remodeling: it enhances collagen-induced platelet activation^51^, is associated with abnormal myelopoiesis^52^, and is strongly implicated in COPD and emphysema^53,54^. In parallel, coagulation pathways were downregulated (**Fig. 7F**), with decreased expression of GP1bα, GP1bβ, and GPV (**Fig. 7G, Supplemental Table 2**). In summary, *in vivo* IL-33 stimulation preserves thrombus formation but remodels the platelet proteome toward inflammation and adhesion.

**Figure 7.**
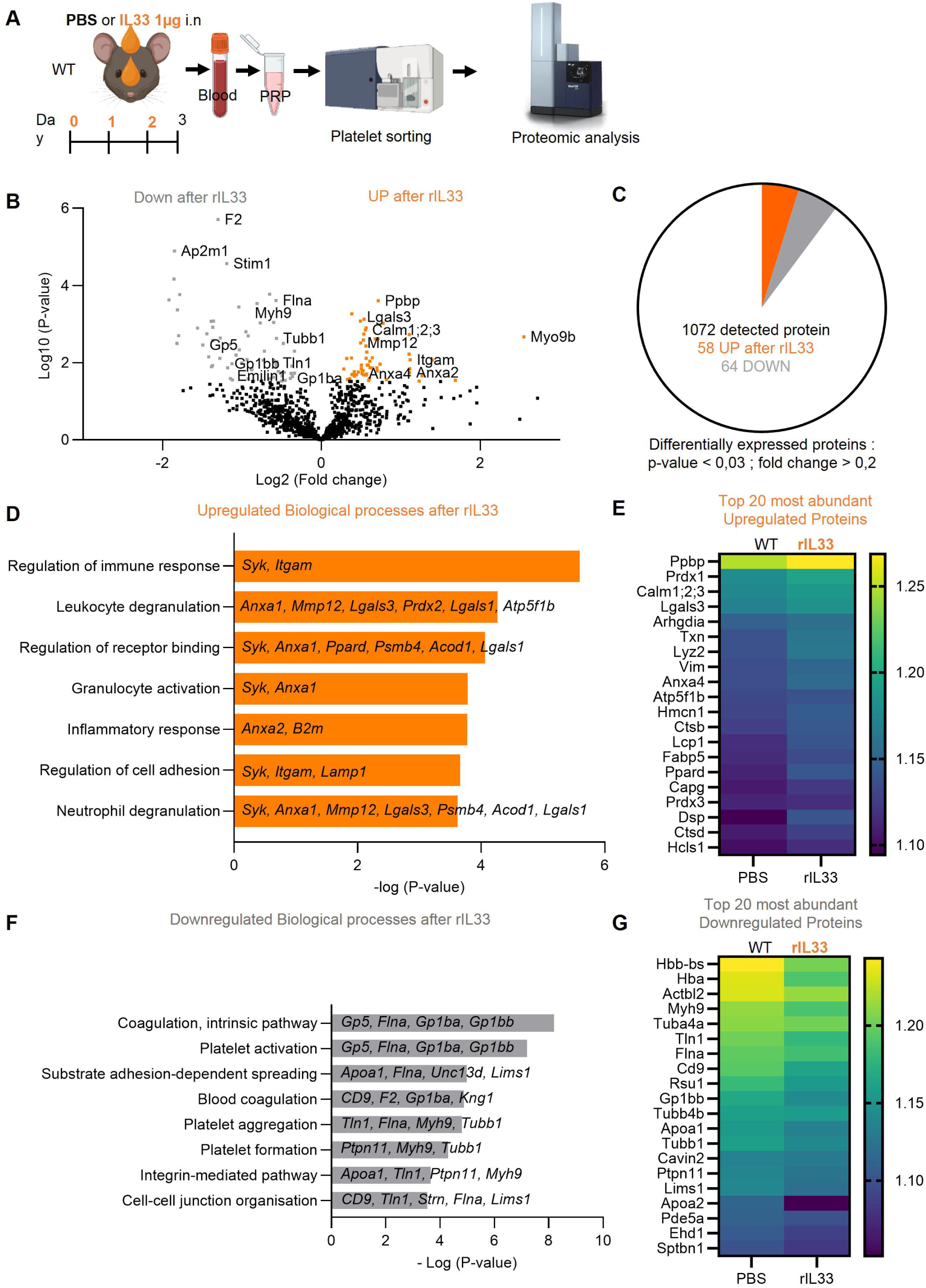
IL-33 *in vivo* treatment alters platelet proteome, modulating inflammatory and coagulation-related pathways (A) Proteomic profiling of sorted platelets from WT mice treated with PBS or rIL-33. (B) Volcano plot comparing rIL-33h and PBS treated mice platelet proteomes. The x-axis shows log fold change (IL-33KO/WT), and the y-axis shows –log(p-value). Proteins upregulated in IL-33 treated mice platelets are highlighted in orange; downregulated proteins in grey. (C) Number of proteins detected and significantly altered (p < 0.03, fold change > 0.2). (D, F) Gene Ontology (GO) analysis of enriched biological processes among upregulated (D) and downregulated (F) proteins (https://geneontology.org). (E, G) Heatmaps of the top 20 most abondant upregulated (E) and downregulated (G) proteins. Data represent n = 10 mice per group, collected from two independent experiments.

## Discussion

Our study uncovers a novel role for IL-33 in platelet biology. We demonstrate that endogenous IL-33 protein is not expressed by circulating platelets, but it indirectly alters their proteome and function. Although a previous study detected IL-33 in platelets, our analyses using IL-33KO mice indicate that the detected signal was probably non-specific. While platelets can internalize circulating proteins, we found no evidence of IL-33 uptake in platelets incubated with recombinant IL-33. Whether IL-33 internalization contributes to platelet function in inflammation remains to be determined.

Interestingly, platelets from IL-33KO mice exhibited functional changes even under homeostatic conditions. Specifically, platelets from IL-33KO mice showed impaired adhesion to matrix proteins (fibrinogen, podoplanin, laminin, and collagen) under both static and flow conditions. However, aggregation and degranulation remained intact, suggesting that IL-33 influences early platelet activation steps, particularly those involving interactions with the endothelium and ECM. These results align with previous findings showing that platelets from IL-33KO mice display reduced eosinophil and neutrophil recruitment in models of pulmonary^15^ and intestinal inflammation^12^. Defective interactions with ECM may also play a key role in megakaryocyte localization and platelet biogenesis, as recently highlighted by the involvement of the Filamin A/GPIb pathway^55^ and the laminin-rich matrix cage surrounding megakaryocytes^56^.

Proteomic analyses further confirmed the role of IL-33 in platelet adhesion. Platelets from IL-33KO mice exhibited downregulation of proteins involved in cytoskeletal remodeling and ECM interaction, including Emilin1, Multimerin-1, Filamin A, and Myh9, all of which are essential for platelet adhesion, migration or thrombus formation ^57–59^. These alterations, along with impaired integrin activation provide a mechanistic explanation for the adhesion defects observed. Additionally, IL-33 may influence platelet migration during inflammation, a Myh9 dependant process known as haptotaxis, where platelets extend lamellipodia and migrate along ECM gradients^59^.

*In vivo*, IL-33 stimulation induced platelet activation, as evidenced by increased plasma sCD40L and degranulation markers. IL-33 also induced a proteomic shift toward an inflammatory phenotype, with upregulation of granulocyte-recruiting proteins like PPBP and downregulation of coagulation-related proteins. This reflects alterations observed in platelets from patients with severe allergies, characterized by elevated levels of MMRN1, CD40L and PPBP^16^ as well as in newly generated platelets following acute thrombocytopenia, which display impaired haemostatic functions and increased expression of immune proteins (TLRs, CD40L, CD63…)^60^.

IL-33 also appears to promote thrombopoiesis. In IL-33 stimulated mice, the presence of large circulating platelets, indicative of de novo production, and the fragmentation of proplatelets within lung capillaries suggest enhanced thrombopoietic activity. The stability of platelet counts likely reflects a balance between increased platelet activation and consumption, and a compensatory rise in production. Notably, the lungs are an efficient site for platelet biogenesis under stress^44^, able to fragment a mature megakaryocyte into platelets in under an hour, much faster than in bone marrow^47^. IL-33 may thus support platelet homeostasis by promoting efficient thrombopoiesis during inflammation.

Mechanistically, IL-33 does not act directly on platelets, as they lack the ST2 receptor. Instead, it likely operates through intermediary cells that express ST2. Flow cytometry and RNA-seq identified ST2 expression on specific hematopoietic and megakaryocyte progenitor subsets in mouse and human bone marrow and lungs. These include megakaryocyte-biased hematopoietic stem cells, especially those involved in non-canonical thrombopoiesis^32,61^. IL-33 may influence platelet production by affecting the maturation of these progenitors. Moreover, the existence of distinct lung-resident and immune-associated megakaryocyte populations suggests that IL-33 selectively engages non-canonical, rapid-response pathways of platelet production. *In vivo* lineage-tracing experiments will be required to determine whether these ST2^+^ progenitor populations are connected to mature megakaryocytes and platelets.

This is consistent with evidence showing ST2 enrichment in non-canonical progenitors in aged mice, as well as reduced expression of immune-related proteins in lung MKs following ST2 blockade^23^. With age, the proportion of megakaryocyte-biased HSCs, populations expressing the IL-33 receptor ST2, increases, suggesting that IL-33 responsiveness may be enhanced in the aged hematopoietic system. IL-33 is constitutively expressed by stromal cells and released upon cellular stress or damage. Since tissue injury and chronic low-grade inflammation rise with age, extracellular IL-33 levels are likely elevated, consistent with observations in bone marrow fluid from aged mice^62^. Together, these findings suggest that age-associated increases in IL-33 release and signalling within the hematopoietic niche may promote the expansion of megakaryocyte-biased and alternate HSC populations. Further studies should test this hypothesis directly.

IL-33 may also indirectly promote megakaryopoiesis via group 2 innate lymphoid cells (ILC2s), which produce IL-6, a known driver of megakaryocyte maturation. IL-33 overexpression increases platelet counts in an IL-6–dependent manner^11^. IL-33-induced IL-6 production from ILC2s is associated with myeloid bias under stress or aging conditions^62^. These indirect pathways support IL-33’s role in linking immune activation with hematopoietic differentiation. Future studies employing MK-, HSC-, or ILC2-specific ST2 knockout mice will be essential to directly assess the contribution of IL-33 signalling within these cell populations to platelet phenotypes.

IL-33 deficiency and acute IL-33 stimulation have distinct effects on platelet proteome and function. IL-33KO mice reveal the physiological role of IL-33 expressed by stromal cells from birth, including during development, while recombinant IL-33 treatment models an acute response to inflammation or tissue injury. This difference likely reflects the involvement of distinct target cells whose responses depend on spatial and temporal context, particularly during the perinatal period^63,64^ or hematopoietic stress and ageing^62^, when IL-33 signalling and ST2⁺ cells may play more prominent roles.

In summary, our findings identify IL-33 as a key regulator of platelet production and function. Rather than acting directly on platelets, IL-33 likely shapes them through effects on megakaryocyte differentiation and inflammatory signalling. IL-33 affects platelet adhesion, progenitor differentiation and release, and promotes an inflammatory platelet phenotype, all of which may be clinically relevant in thrombo-inflammatory diseases. These results position IL-33 as a potential therapeutic target for modulating platelet activity and production in inflammation, infection, or aging. Future research should clarify the intermediary cells involved, its role in platelet immune functions, and determine its impact on megakaryocyte differentiation during physiological and pathological stress. We also acknowledge the translational limitations of our mouse-focused study and emphasize that further validation in human hematopoietic progenitors, megakaryocytes and platelets will be necessary to confirm the clinical relevance of our observations.

## Acknowledgments

We acknowledge the proteomics (ProteoToul), cytometry (TRI-IPBS), imaging (TRI-IPBS) and animal facilities (Anexplo-IPBS) for technical support. TRI-IPBS Cytometry and TRI-IPBS Microscopy (Genotoul-TRI) are member of the national infrastructure France-BioImaging (https://ror.org/01y7vt929) supported by the French National Research Agency (ANR-24-INBS-0005 FBI BIOGEN). This work was supported by the French ministry of Research, ANR JCJC (Grant #ANR-20-CE14-0045-01), Fonds de Recherche en Santé Respiratoire - Fondation du Souffle (Grant #198942). LG is supported by a PhD fellowship from SFH (Société Française d’Hématologie). Figures were created with Biorender.com. We acknowledge Andrew McKenzie (MRC, Cambridge) for kindly providing the IL-33^cit/cit^ mice.

## Authorship Contributions

L.G. conducted the experiments and drafted the manuscript. S.R. produced the recombinant IL-33 protein. A.G.P. and E.V. carried out the proteomic experiments and analysis. S.S. performed the thrombus formation assays under flow conditions and experiments involving aged mice. J.-P.G. provided financial support, and contributed to conceptual development. E.L. supervised the research, designed and performed experiments, and wrote the manuscript. All authors discussed the results and contributed to the final manuscript.

## References

1. Nicolai L, Pekayvaz K, Massberg S. Platelets: Orchestrators of immunity in host defense and beyond. Immunity. 2024;57(5):957–972. doi:10.1016/j.immuni.2024.04.008

2. Li JL, Zarbock A, Hidalgo A. Platelets as autonomous drones for hemostatic and immune surveillance. Journal of Experimental Medicine. 2017;214(8):2193–2204. doi:10.1084/jem.20170879

3. Middleton EA, Weyrich AS, Zimmerman GA. Platelets in Pulmonary Immune Responses and Inflammatory Lung Diseases. Physiological Reviews. 2016;96(4):1211–1259. doi:10.1152/physrev.00038.2015

4. Cayrol C, Girard JP. Interleukin-33 (IL-33): A critical review of its biology and the mechanisms involved in its release as a potent extracellular cytokine. Cytokine. 2022;156:155891. doi:10.1016/j.cyto.2022.155891

5. Moussion C, Ortega N, Girard JP. The IL-1-Like Cytokine IL-33 Is Constitutively Expressed in the Nucleus of Endothelial Cells and Epithelial Cells In Vivo: A Novel ‘Alarmin’? Unutmaz D, ed. PLoS ONE. 2008;3(10):e3331. doi:10.1371/journal.pone.0003331

6. Pichery M, Mirey E, Mercier P, et al. Endogenous IL-33 Is Highly Expressed in Mouse Epithelial Barrier Tissues, Lymphoid Organs, Brain, Embryos, and Inflamed Tissues: In Situ Analysis Using a Novel *Il-33–LacZ* Gene Trap Reporter Strain. The Journal of Immunology. 2012;188(7):3488–3495. doi:10.4049/jimmunol.1101977

7. Cayrol C, Girard JP. Interleukin-33 (IL-33): A nuclear cytokine from the IL-1 family. Immunol Rev. 2018;281(1):154–168. doi:10.1111/imr.12619

8. Alt C, Yuan S, Wu F, et al. Long-Acting IL-33 Mobilizes High-Quality Hematopoietic Stem and Progenitor Cells More Efficiently Than Granulocyte Colony-Stimulating Factor or AMD3100. Biol Blood Marrow Transplant. 2019;25(8):1475–1485. doi:10.1016/j.bbmt.2019.05.030

9. Kim J, Kim W, Le HT, et al. IL-33-induced hematopoietic stem and progenitor cell mobilization depends upon CCR2. J Immunol. 2014;193(7):3792–3802. doi:10.4049/jimmunol.1400176

10. Swann JW, Koneva LA, Regan-Komito D, Sansom SN, Powrie F, Griseri T. IL-33 promotes anemia during chronic inflammation by inhibiting differentiation of erythroid progenitors. J Exp Med. 2020;217(9):e20200164. doi:10.1084/jem.20200164

11. Talabot-Ayer D, Martin P, Vesin C, et al. Severe Neutrophil-Dominated Inflammation and Enhanced Myelopoiesis in IL-33–Overexpressing CMV/IL33 Mice. The Journal of Immunology. 2015;194(2):750–760. doi:10.4049/jimmunol.1402057

12. Chen Z, Luo J, Li J, et al. Intestinal IL-33 promotes platelet activity for neutrophil recruitment during acute inflammation. Blood. 2022;139(12):1878–1891. doi:10.1182/blood.2021013474

13. Takeda T, Unno H, Morita H, et al. Platelets constitutively express IL-33 protein and modulate eosinophilic airway inflammation. J Allergy Clin Immunol. 2016;138(5):1395–1403.e6. doi:10.1016/j.jaci.2016.01.032

14. Liu T, Barrett NA, Kanaoka Y, et al. Cysteinyl leukotriene receptor 2 drives lung immunopathology through a platelet and high mobility box 1-dependent mechanism. Mucosal Immunology. 2019;12(3):679–690. doi:10.1038/s41385-019-0134-8

15. Zeiler M, Moser M, Mann M. Copy Number Analysis of the Murine Platelet Proteome Spanning the Complete Abundance Range. Molecular & Cellular Proteomics. 2014;13(12):3435–3445. doi:10.1074/mcp.m114.038513

16. Pablo-Torres C, Izquierdo E, Tan TJ, et al. Deciphering the role of platelets in severe allergy by an integrative omics approach. *A*llergy. 2023;78(5):1319–1332. doi:10.1111/all.15621

17. Wang H, He J, Xu C, et al. Decoding Human Megakaryocyte Development. Cell Stem Cell. 2021;28(3):535–549.e8. doi:10.1016/j.stem.2020.11.006

18. Sun S, Jin C, Si J, et al. Single-cell analysis of ploidy and the transcriptome reveals functional and spatial divergency in murine megakaryopoiesis. Blood. 2021;138(14):1211–1224. doi:10.1182/blood.2021010697

19. Valet C, Magnen M, Qiu L, et al. Sepsis promotes splenic production of a protective platelet pool with high CD40 ligand expression. Journal of Clinical Investigation. 2022;132(7). doi:10.1172/jci153920

20. Wang J, Xie J, Wang D, et al. CXCR4high megakaryocytes regulate host-defense immunity against bacterial pathogens. eLife. 2022;11:e78662. doi:10.7554/eLife.78662

21. Lefrançais E, Ortiz-Muñoz G, Caudrillier A, et al. The lung is a site of platelet biogenesis and a reservoir for haematopoietic progenitors. Nature. 2017;544(7648):105–109. doi:10.1038/nature21706

22. Yeung AK, Villacorta-Martin C, Hon S, Rock JR, Murphy GJ. Lung megakaryocytes display distinct transcriptional and phenotypic properties. Blood Advances. 2020;4(24):6204–6217. doi:10.1182/bloodadvances.2020002843

23. Pariser DN, Hilt ZT, Ture SK, et al. Lung megakaryocytes are immune modulatory cells. Journal of Clinical Investigation. 2021;131(1). doi:10.1172/jci137377

24. Looney MR, Thornton EE, Sen D, Lamm WJ, Glenny RW, Krummel MF. Stabilized imaging of immune surveillance in the mouse lung. Nat Methods. 2011;8(1):91–96. doi:10.1038/nmeth.1543

25. Headley MB, Bins A, Nip A, et al. Visualization of immediate immune responses to pioneer metastatic cells in the lung. Nature. 2016;531(7595):513–517. doi:10.1038/nature16985

26. Luigi Grassi, Osagie G. Izuogu, Natasha A.N. Jorge, et al. Cell type-specific novel long non-coding RNA and circular RNA in the BLUEPRINT hematopoietic transcriptomes atlas. haematol. 2020;106(10):2613–2623. doi:10.3324/haematol.2019.238147

27. Burkhart JM, Vaudel M, Gambaryan S, et al. The first comprehensive and quantitative analysis of human platelet protein composition allows the comparative analysis of structural and functional pathways. Blood. 2012;120(15):e73–e82. doi:10.1182/blood-2012-04-416594

28. Lefrançais E, Roga S, Gautier V, et al. IL-33 is processed into mature bioactive forms by neutrophil elastase and cathepsin G. Proc Natl Acad Sci U S A. 2012;109(5):1673–1678. doi:10.1073/pnas.1115884109

29. Cayrol C, Duval A, Schmitt P, et al. Environmental allergens induce allergic inflammation through proteolytic maturation of IL-33. Nat Immunol. 2018;19(4):375–385. doi:10.1038/s41590-018-0067-5

30. Schmitt P, Duval A, Camus M, et al. TL1A is an epithelial alarmin that cooperates with IL-33 for initiation of allergic airway inflammation. J Exp Med. 2024;221(6):e20231236. doi:10.1084/jem.20231236

31. Hardman CS, Panova V, McKenzie ANJ. IL-33 citrine reporter mice reveal the temporal and spatial expression of IL-33 during allergic lung inflammation. Eur J Immunol. 2013;43(2):488–498. doi:10.1002/eji.201242863

32. Livada AC, McGrath KE, Malloy MW, et al. Long-lived lung megakaryocytes contribute to platelet recovery in thrombocytopenia models. Journal of Clinical Investigation. 2024;134(22). doi:10.1172/jci181111

33. Noetzli LJ, French SL, Machlus KR. New Insights Into the Differentiation of Megakaryocytes From Hematopoietic Progenitors. ATVB. 2019;39(7):1288–1300. doi:10.1161/atvbaha.119.312129

34. Gekas C, Graf T. CD41 expression marks myeloid-biased adult hematopoietic stem cells and increases with age. Blood. 2013;121(22):4463–4472. doi:10.1182/blood-2012-09-457929

35. Manso BA, Medina P, Smith-Berdan S, et al. A rare HSC-derived megakaryocyte progenitor accumulates via enhanced survival and contributes to exacerbated thrombopoiesis upon aging. Cold Spring Harbor Laboratory. Preprint posted online 5 November 2024. doi:10.1101/2024.11.04.621964

36. Morcos MNF, Li C, Munz CM, et al. Fate mapping of hematopoietic stem cells reveals two pathways of native thrombopoiesis. Nat Commun. 2022;13(1). doi:10.1038/s41467-022-31914-z

37. Poscablo DM, Worthington AK, Smith-Berdan S, et al. An age-progressive platelet differentiation path from hematopoietic stem cells causes exacerbated thrombosis. Cell. 2024;187(12):3090–3107.e21. doi:10.1016/j.cell.2024.04.018

38. Haas S, Hansson J, Klimmeck D, et al. Inflammation-Induced Emergency Megakaryopoiesis Driven by Hematopoietic Stem Cell-like Megakaryocyte Progenitors. Cell Stem Cell. 2015;17(4):422–434. doi:10.1016/j.stem.2015.07.007

39. Rommel MGE, Walz L, Fotopoulou F, et al. Influenza A virus infection instructs hematopoiesis to megakaryocyte-lineage output. Cell Reports. 2022;41(1):111447. doi:10.1016/j.celrep.2022.111447

40. Psaila B, Wang G, Rodriguez-Meira A, et al. Single-Cell Analyses Reveal Megakaryocyte-Biased Hematopoiesis in Myelofibrosis and Identify Mutant Clone-Specific Targets. Molecular Cell. 2020;78(3):477–492.e8. doi:10.1016/j.molcel.2020.04.008

41. Ho YH, Del Toro R, Rivera-Torres J, et al. Remodeling of Bone Marrow Hematopoietic Stem Cell Niches Promotes Myeloid Cell Expansion during Premature or Physiological Aging. Cell Stem Cell. 2019;25(3):407–418.e6. doi:10.1016/j.stem.2019.06.007

42. Mende N, Bastos HP, Santoro A, et al. Unique molecular and functional features of extramedullary hematopoietic stem and progenitor cell reservoirs in humans. Blood. 2022;139(23):3387–3401. doi:10.1182/blood.2021013450

43. Li R. Platelet size matters. Blood. 2024;143(4):298–300. doi:10.1182/blood.2023023057

44. Gelon L, Fromont L, Lefrançais E. Occurrence and role of lung megakaryocytes in infection and inflammation. Front Immunol. 2022;13. doi:10.3389/fimmu.2022.1029223

45. Lefrançais E, Looney MR. Platelet Biogenesis in the Lung Circulation. Physiology (Bethesda). 2019;34(6):392–401. doi:10.1152/physiol.00017.2019

46. Zhao X, Alibhai D, Walsh TG, et al. Highly efficient platelet generation in lung vasculature reproduced by microfluidics. Nat Commun. 2023;14(1):4026. doi:10.1038/s41467-023-39598-9

47. Gaertner F, Ishikawa-Ankerhold H, Stutte S, et al. Plasmacytoid dendritic cells control homeostasis of megakaryopoiesis. Nature. 2024;631(8021):645–653. doi:10.1038/s41586-024-07671-y

48. Graca FA, Stephan A, Minden-Birkenmaier BA, et al. Platelet-derived chemokines promote skeletal muscle regeneration by guiding neutrophil recruitment to injured muscles. Nat Commun. 2023;14(1):2900. doi:10.1038/s41467-023-38624-0

49. Mammadova-Bach E, Gil-Pulido J, Sarukhanyan E, et al. Platelet glycoprotein VI promotes metastasis through interaction with cancer cell-derived galectin-3. Blood. 2020;135(14):1146–1160. doi:10.1182/blood.2019002649

50. Kahner BN, Dorsam RT, Mada SR, et al. Hematopoietic lineage cell specific protein 1 (HS1) is a functionally important signaling molecule in platelet activation. Blood. 2007;110(7):2449–2456. doi:10.1182/blood-2006-11-056069

51. Wang J, Ye Y, Wei G, et al. Matrix metalloproteinase12 facilitated platelet activation by shedding carcinoembryonic antigen related cell adhesion molecule1. Biochem Biophys Res Commun. 2017;486(4):1103–1109. doi:10.1016/j.bbrc.2017.04.001

52. Qu P, Yan C, Du H. Matrix metalloproteinase 12 overexpression in myeloid lineage cells plays a key role in modulating myelopoiesis, immune suppression, and lung tumorigenesis. Blood. 2011;117(17):4476–4489. doi:10.1182/blood-2010-07-298380

53. Hunninghake GM, Cho MH, Tesfaigzi Y, et al. MMP12, lung function, and COPD in high-risk populations. N Engl J Med. 2009;361(27):2599–2608. doi:10.1056/NEJMoa0904006

54. Hautamaki RD, Kobayashi DK, Senior RM, Shapiro SD. Requirement for macrophage elastase for cigarette smoke-induced emphysema in mice. Science. 1997;277(5334):2002–2004. doi:10.1126/science.277.5334.2002

55. Ellis ML, Terreaux A, Alwis I, et al. GPIbα-filamin A interaction regulates megakaryocyte localization and budding during platelet biogenesis. Blood. 2024;143(4):342–356. doi:10.1182/blood.2023021292

56. Masson C, Scandola C, Rinckel JY, et al. Megakaryocytes assemble a three-dimensional cage of extracellular matrix that controls their maturation and anchoring to the vascular niche. Preprint posted online 10 February 2025. doi:10.7554/eLife.104963.1

57. Leatherdale A, Parker D, Tasneem S, et al. Multimerin 1 supports platelet function in vivo and binds to specific GPAGPOGPX motifs in fibrillar collagens that enhance platelet adhesion. J Thromb Haemost. 2021;19(2):547–561. doi:10.1111/jth.15171

58. Rosa JP, Raslova H, Bryckaert M. Filamin A: key actor in platelet biology. Blood. 2019;134(16):1279–1288. doi:10.1182/blood.2019000014

59. Gaertner F, Ahmad Z, Rosenberger G, et al. Migrating Platelets Are Mechano-scavengers that Collect and Bundle Bacteria. Cell. 2017;171(6):1368–1382.e23. doi:10.1016/j.cell.2017.11.001

60. Pirabe A, Schrottmaier WC, Mehic D, et al. Impaired hemostatic and immune functions of platelets after acute thrombocytopenia. Journal of Thrombosis and Haemostasis. 2025;23(3):1052–1065. doi:10.1016/j.jtha.2024.11.029

61. Li JJ, Liu J, Li YE, et al. Differentiation route determines the functional outputs of adult megakaryopoiesis. Immunity. 2024;57(3):478–494.e6. doi:10.1016/j.immuni.2024.02.006

62. Naef P, Jaeger-Ruckstuhl CA, Schnüriger N, et al. IL-33/ST2 signaling in ILC2s drives exhaustion and myeloid skewing of HSCs in response to hematopoietic stress and aging. iScience. 2025;28(5):112378. doi:10.1016/j.isci.2025.112378

63. de Kleer IM, Kool M, de Bruijn MJW, et al. Perinatal Activation of the Interleukin-33 Pathway Promotes Type 2 Immunity in the Developing Lung. Immunity. 2016;45(6):1285–1298. doi:10.1016/j.immuni.2016.10.031

64. Saluzzo S, Gorki AD, Rana BMJ, et al. First-Breath-Induced Type 2 Pathways Shape the Lung Immune Environment. Cell Rep. 2017;18(8):1893–1905. doi:10.1016/j.celrep.2017.01.071

