## Supplementary material for "The alarmin Interleukin-33 modulates platelet proteome, function and biogenesis": Suplemental Data

**Supplemental Methods**

***Mice***

Female wild-type WT and IL-33-deficient (IL-33 KO) mice on a C57BL/6J background (littermates, 2–6 months) were used and previously described[1]. $Il33^{Cit/Cit}$ reporter mice were kindly provided by A. McKenzie (MRC, Cambridge, UK)[2]. PF4-mTmG and PF4-nTnG reporter mice were generated by crossing PF4-Cre mice with *mTmG* or *nTnG* reporter strains (Jackson Laboratory). Ageing experiments compared 8-week-old with 2-year-old mice. Animals were randomly assigned to experimental groups using littermates to minimize genetic and environmental variability. When indicated in the figure legends, investigators were blinded to genotype and/or treatment during data collection and analysis. Group sizes were determined based on previous studies with comparable experimental endpoints to ensure adequate statistical power (typically $\geq 0.8$, $\alpha = 0.05$). The number of biological and technical replicates is provided in the figure legends. Only female mice were included in this study, and sex-based differences were not evaluated. All animal experiments were conducted in accordance with European and French ethical regulations and approved by the institutional ethics committee.

***IL-33 recombinant protein production and purification***

cDNAs encoding human $IL\text{-}33_{112–270}$ WT were amplified by PCR with human *IL33* cDNA (NM_033439.3) as the template. For expression in Escherichia coli, cDNAs encoding mature form IL-33(112–270) WT were subcloned into expression vector pET-15b (Novagen). Recombinant proteins were produced in E. coli BL21pLysS (Novagen) and purified using Ni-NTA affinity agarose column (Qiagen). His tag was removed by cleavage with thrombin, and proteins were further purified by gel filtration (FPLC; GE Healthcare). Recombinant Il-33 proteins were stored in 50mM phosphate pH 7,5 buffer in the presence of 5mM DTT, to increase protein stability during long term storage, because IL-33 is sensitive to inactivation by oxidation [3]. Endotoxin levels were <0.01 EU/µg of protein, as determined by the Limulus amebocyte lysate QCL-1000 method (Lonza).

***Administration of IL-33 in vivo***

Mice were anesthetized, followed by 3 daily i.n. administration of 50 µl of recombinant human IL-33 ($rIL\text{-}33_{112\text{-}270}$, 1 µg; house-made), in PBS. This protocol was previously validated for detecting IL-33–specific biological activity activity[4,5]. Blood and bone marrow were collected 24 h after the last treatment with IL-33 for flow cytometry, hematoanalysis, and ELISA.

*Antibodies and reagents*

Antibodies and reagents are provided in Supplemental table 3 and 4.

*Platelet-rich plasma (PRP) preparation*

Intracardiac blood was collected using a 26G needle coated with 150 µL ACD. 500 µL was mixed with 1 mL PIPES and 0.5 µL 1 µM $PGI_2$, then centrifuged (70g, 20 °C, 10 min, no brake). The supernatant (~1 mL) was centrifuged (600g, 20 °C, 7 min), and the pellet resuspended in 1 mL Tyrode's buffer (136 mM NaCl, 2,68 mM KCl, 1mM $MgCl_2$, 1 mM $CaCl_2$, 11,9 mM $NaHCO_3$, 0.4mM $NaH_2PO_4$, 0.8mM glucose, 0.35% BSA and 2,5mM HEPES, pH 7.4).

*Megakaryocyte Enrichment*

500 µL of bone marrow suspension was layered on a BSA gradient (1.5 mL PBS/3% BSA over 1.5 mL PBS/1.5% BSA). After 40 min at RT, the lower phase was collected, centrifuged (400g, 5 min, 20 °C), and resuspended in 1 mL Tyrode's buffer with inhibitors (Apyrase 2U/mL, Heparin 10U/mL, PGI2 1µM).

*IL-33 expression analysis by flow cytometry*

Reagents and antibodies are listed in Supplemental Tables 3–4. PRP was prepared as described and stained (15 min, RT) with CD41-BV786, CD45-AF700, and Ter119-APC. After centrifugation (600g, 7 min, 20 °C), platelets were fixed/permeabilized (FOXP3 kit), washed, and incubated with anti-IL-33 (20 min,) and Cy3 anti-goat IgG (20 min) was added. After a final wash (600g, 7 min, 20 °C); platelets were resuspended in 200 µL Tyrode's buffer.

*Platelet isolation for proteomic analysis*

Platelet-rich plasma was stained with CD41-BV786, CD45-AF700, and Ter119-APC and sorted on an ARIA cytometer. Absence of platelet activation during the process was assessed pre/post-sorting using P-selectin staining.

*Platelet sample preparation for mass spectrometry and analysis*

Purified platelets (min $5.10^6$) were centrifuged (600g, 7 min, 20 °C), and prepared for proteomic analysis. Platelets were lysed in a solution of 5% sodium dodecyl sulfate (SDS) (Invitrogen, 15553-035) and 50 mM triethylammonium bicarbonate, pH 8.5. Protein disulfide bridges and cysteine residues were reduced and alkylated by incubation with 10mMTris (2-carboxyethyl) phosphine hydrochloride (NeoBiotech, 45-00029-10g) and 40mM 2-chloroacetamide (Aldrich, C0267-100G), 5min at pH 8.5 and 95°C. Proteins were digested on S-trap Micro devices (Protifi) using the manufacturer's protocol. Briefly, samples were acidified by addition of phosphoric acid (2.5% final concentration), and proteins were precipitated by addition of 6 volumes of 90% methanol in 100mM TEAB, pH7.5. Samples were applied on the S-Trap Micro devices and precipitated proteins were washed 6 times with 90% methanol, 100mM TEAB, pH7.5. Proteins were then digested by incubation with trypsin (catalog no. V5117, Promega) at 1% (trypsin/protein) and 37°C overnight. The resulting peptides were succesively eluted with 40µl of 50mM ammonium bicarbonate, 40µl of 0.2% formic acid and 40µl of 50% acetonitrile, 0.2% formic acid. and speed-vacuum dried. After resuspension in 0.1% formic acid, they were desalted on Affinisep C18 T1 Tips with 0.1% formic acid, eluted with 40% acetonitrile, 0.1% formic acid and then with 70% acetonitrile, 0.1% formic acid, and finally dried again.

Peptides were analyzed by nanoscale liquid chromatography using an UltiMate 3000 system (NCS-3500RS Nano/Cap System, Thermo Scientific) coupled to a timsTOF SCP mass spectrometer (Bruker Daltonics). Peptides were separated on a C18 Aurora column (25 cm x 75 µm ID, IonOpticks) using a gradient ramping from 2% to 20% of solvent B in 30 min, then to 37% of solvent B in 3 min and to 85% of solvent B in 2min (solvent A: 0.1% formic acid in H2O; solvent B 0.1% formic acid in acetonitrile), with a flow rate of 150 nl/min. MS acquisition was performed in DIA-PASEF mode on the precursor mass range [400-1000] m/z and ion mobility 1/K0 [0.64-1.37]. The acquisition scheme was composed of 8 consecutive TIMS ramps using an accumulation time of 100 ms, with 3 MS/MS acquisition windows of 25 Th for each of them. The resulting cycle time was 0.96 s. The collision energy was ramped linearly as a function of the ion mobility from 59 eV at 1/K0=1.6 Vs.cm−2 to 20 eV at 1/K0=0.6Vs.cm−2.

The raw data was searched and quantified with the DIA-NN software (version 1.8.1), using a predicted library generated by the software from the Uniprot mouse reference proteome (download April 2023). Inclusion criteria for precursor ions hits were a peptide length range set at 7–30, and a precursor charge range set at 2–3. Validation of the identifications was performed using the target-decoy approach implemented in DIA-NN setting the final false discovery rate at 1% both at precursor ion level and protein level. With these settings, an average proteomic depth of 1428 proteins identified per sample was obtained for the experiment comparing the proteome of platelets purified by FACS from IL-33KO or WT mice. For the experiment comparing the proteome of platelets from WT mice either non stimulated or treated with recombinant IL-33, around 882 proteins were identified in average across the samples. For each experiment, the "global normalization" option was selected in DIA-NN to correct for different amounts of input material and sample losses. Protein intensities reported in the "report.pg_matrix" output from DIA-NN were used for calculation of fold changes and statistical analysis using a Student t-test calculated between the two conditions compared (n=5 biological replicates in each of the two groups for the KO vs WT experiment; n=8 or 9 biological replicates in the two respective groups for the Non-treated vs IL33-treated experiment). To select differentially abundant candidate proteins, threshold values of p-val < 0.03 and absolute log2(fold change) > 0.2 were applied. The mass spectrometry proteomics data have been deposited to the ProteomeXchange Consortium via the PRIDE [6] partner repository with the dataset identifier PXD068068.

***Blood platelet analysis by flow cytometry***

A 50 µL blood sample was collected from the mandible using an ACD-coated Minivette and diluted 1:20 in PBS$^{-/-}$ with 5 mM EDTA. Samples were stained for 15 min at room temperature in the dark with antibodies listed in Supplemental Table 3. For quantification of the immature platelet fraction, platelets were stained with 1 µM thiazole orange. For platelet quantification, count beads were added (1:1). Acquisition was performed on a Fortessa X20 or Northern Lights flow cytometer. For platelet analysis, the FSC voltage and threshold were adapted using size-calibrated beads to ensure accurate detection of small events (Megamix-Plus, Stago).

***Hematopoietic stem and progenitor cell analysis by flow cytometry***

Tibias and femurs from both legs were removed. Bone marrow cells were flushed with PBS with 5 mM EDTA before 100-µm cell strainer filtration and red blood cell lysis. Bone marrow cells were resuspended in FACS buffer, and Fc receptors were blocked using CD16/32 antibody along with Viability Red Dye. Cells were subsequently stained with a lineage (Lin) antibody cocktail conjugated to FITC, including CD19, CD11b, CD4, CD3, CD45R, CD11c, NK1.1, Ter119, Gr1, FcεRI. Additional staining included antibodies against CD45, CD41, c-Kit, Sca-1, CD48, CD150, ST2, CD127, and CD90.2. Hematopoietic stem and progenitor cell (HSPC) populations were defined as follows: after exclusion of ILCs (Lin$^-$ Cd45$^+$ CD90.2$^+$ CD127$^+$), the LSK compartment was defined as Lin$^-$ CD45$^+$ Sca-1$^+$ c-Kit$^+$, and the LK compartment as Lin$^-$ CD45$^+$ Sca-1$^-$ c-Kit$^+$. Megakaryocyte progenitors (MkPs) were identified as LK CD150$^+$ CD41$^+$, short-term HSCs (ST-HSCs) as LSK CD48$^-$ CD150$^-$, and long-term HSCs (LT-HSCs) as LSK CD48$^-$ CD150$^+$. MK-biased LT-HSCs were defined as CD41$^+$ LT-HSCs, and MK-biased MkPs as CD48$^-$ MkPs. Antibodies are listed in Supplemental Table 3. Samples were analysed on a Northern Light spectral cytometer (Cytek). Instrument were set using unstained and single-stained controls, and unmixing was applied automatically based on singlecolor controls. Doublets were excluded by gating on FSC-A versus FSC-H and SSC-A versus SSC-H. Analysis of flow cytometry data was performed using Flowjo (Treestar).

##### *Washed platelet preparation and Aggregation*

Intracardiac blood was collected with a 1 mL syringe containing 150 µL ACD. Blood was diluted in modified HEPES-Tyrode buffer (pH 6.7) with glucose (5.5 mM) and 0.35% BSA, then centrifuged at 300g, 24°C for 4 min (accel. 7, brake 7) to obtain PRP. After adding 500 nM PGI2, PRP was centrifuged at 1000g, 24°C for 6 min (accel. 7, brake 7). The platelet pellet was resuspended in HEPES-Tyrode buffer (pH 7.3) with glucose (5.5 mM) and 0.02 U/mL apyrase, rested at 37°C for 45 min, and adjusted to $2\times10^8$ platelets/mL. Washed platelet aggregation was measured turbidimetrically in an octochannel aggregometer at 37°C, stirring at 1000 rpm, after stimulation with collagen (0.5 µg/mL), CRP (0.1 or 1 µg/mL), or thrombin (0.1 IU/mL).

##### *Supplemental References*


1. Pichery M, Mirey E, Mercier P, et al. Endogenous IL-33 Is Highly Expressed in Mouse Epithelial Barrier Tissues, Lymphoid Organs, Brain, Embryos, and Inflamed Tissues: In Situ Analysis Using a Novel *Il-33–LacZ* Gene Trap Reporter Strain. *The Journal of Immunology*. 2012;188(7):3488-3495. doi:10.4049/jimmunol.1101977

2. Hardman CS, Panova V, McKenzie ANJ. IL-33 citrine reporter mice reveal the temporal and spatial expression of IL-33 during allergic lung inflammation. *Eur J Immunol*. 2013;43(2):488-498. doi:10.1002/eji.201242863

3. Cohen ES, Scott IC, Majithiya JB, et al. Oxidation of the alarmin IL-33 regulates ST2-dependent inflammation. *Nat Commun*. 2015;6:8327. doi:10.1038/ncomms9327

4. Kondo Y, Yoshimoto T, Yasuda K, et al. Administration of IL-33 induces airway hyperresponsiveness and goblet cell hyperplasia in the lungs in the absence of adaptive immune system. *Int Immunol*. 2008;20(6):791-800. doi:10.1093/intimm/dxn037

5. Oboki K, Ohno T, Kajiwara N, et al. IL-33 is a crucial amplifier of innate rather than acquired immunity. *Proc Natl Acad Sci U S A*. 2010;107(43):18581-18586. doi:10.1073/pnas.1003059107

6. Perez-Riverol Y, Bandla C, Kundu DJ, et al. The PRIDE database at 20 years: 2025 update. *Nucleic Acids Res*. 2025;53(D1):D543-D553. doi:10.1093/nar/gkae1011

7. Molofsky AB, Van Gool F, Liang HE, et al. Interleukin-33 and Interferon-γ Counter-Regulate Group 2 Innate Lymphoid Cell Activation during Immune Perturbation. *Immunity*. 2015;43(1):161-174. doi:10.1016/j.immuni.2015.05.019

8. Ferhat M, Robin A, Giraud S, et al. Endogenous IL-33 Contributes to Kidney Ischemia-Reperfusion Injury as an Alarmin. *J Am Soc Nephrol*. 2018;29(4):1272-1288. doi:10.1681/ASN.2017060650

9. Dahlgren MW, Jones SW, Cautivo KM, et al. Adventitial Stromal Cells Define Group 2 Innate Lymphoid Cell Tissue Niches. *Immunity*. 2019;50(3):707-722.e6. doi:10.1016/j.immuni.2019.02.002

10. Barbier L, Robin A, Sindayigaya R, et al. Endogenous Interleukin-33 Acts as an Alarmin in Liver Ischemia-Reperfusion and Is Associated With Injury After Human Liver Transplantation. *Front Immunol*. 2021;12:744927. doi:10.3389/fimmu.2021.744927

11. Haenuki Y, Matsushita K, Futatsugi-Yumikura S, et al. A critical role of IL-33 in experimental allergic rhinitis. *J Allergy Clin Immunol*. 2012;130(1):184-194.e11. doi:10.1016/j.jaci.2012.02.013

12. Nakanishi W, Yamaguchi S, Matsuda A, et al. IL-33, but not IL-25, is crucial for the development of house dust mite antigen-induced allergic rhinitis. *PLoS One*. 2013;8(10):e78099. doi:10.1371/journal.pone.0078099

13. Sundnes O, Pietka W, Loos T, et al. Epidermal Expression and Regulation of Interleukin-33 during Homeostasis and Inflammation: Strong Species Differences. *J Invest Dermatol*. 2015;135(7):1771-1780. doi:10.1038/jid.2015.85

14. Aparicio-Domingo P, Cannelle H, Buechler MB, et al. Fibroblast-derived IL-33 is dispensable for lymph node homeostasis but critical for CD8 T-cell responses to acute and chronic viral infection. *Eur J Immunol*. 2021;51(1):76-90. doi:10.1002/eji.201948413

15. Gadani SP, Walsh JT, Smirnov I, Zheng J, Kipnis J. The glia-derived alarmin IL-33 orchestrates the immune response and promotes recovery following CNS injury. *Neuron*. 2015;85(4):703-709. doi:10.1016/j.neuron.2015.01.013

16. Vainchtein ID, Chin G, Cho FS, et al. Astrocyte-derived interleukin-33 promotes microglial synapse engulfment and neural circuit development. *Science*. 2018;359(6381):1269-1273. doi:10.1126/science.aal3589

17. Nguyen PT, Dorman LC, Pan S, et al. Microglial Remodeling of the Extracellular Matrix Promotes Synapse Plasticity. *Cell*. 2020;182(2):388-403.e15. doi:10.1016/j.cell.2020.05.050

18. Cayrol C, Duval A, Schmitt P, et al. Environmental allergens induce allergic inflammation through proteolytic maturation of IL-33. *Nat Immunol*. 2018;19(4):375-385. doi:10.1038/s41590-018-0067-5

19. Lefrançais E, Roga S, Gautier V, et al. IL-33 is processed into mature bioactive forms by neutrophil elastase and cathepsin G. *Proc Natl Acad Sci U S A*. 2012;109(5):1673-1678. doi:10.1073/pnas.1115884109

20. Yasuda K, Muto T, Kawagoe T, et al. Contribution of IL-33-activated type II innate lymphoid cells to pulmonary eosinophilia in intestinal nematode-infected mice. *Proc Natl Acad Sci U S A*. 2012;109(9):3451-3456. doi:10.1073/pnas.1201042109

21. Spallanzani RG, Zemmour D, Xiao T, et al. Distinct immunocyte-promoting and adipocyte-generating stromal components coordinate adipose tissue immune and metabolic tenors. *Sci Immunol*. 2019;4(35):eaaw3658. doi:10.1126/sciimmunol.aaw3658

22. Mahlakõiv T, Flamar AL, Johnston LK, et al. Stromal cells maintain immune cell homeostasis in adipose tissue via production of interleukin-33. *Sci Immunol*. 2019;4(35):eaax0416. doi:10.1126/sciimmunol.aax0416

23. Clark JT, Christian DA, Gullicksrud JA, et al. IL-33 promotes innate lymphoid cell-dependent IFN-γ production required for innate immunity to Toxoplasma gondii. *Elife*. 2021;10:e65614. doi:10.7554/eLife.65614

#### Supplemental Tables

##### Supplemental Table 1. Techniques used for assessing IL-33 expression

| Technique | Positive control | Negative control | Ref * | Result in platelets | Result in megakaryocytes |
|---|---|---|---|---|---|
| **Immunofluorescence (IF) -** Goat Polyclonal anti-mouse IL33 AF3626 | ✅ IL-33$^+$ epithelial cells, endothelial cells or stromal cells in various tissues, or IL-33+ astrocytes, ologodendrocytes in the brain (nuclear staining) | No signal in IL-33 KO mice | 1,6–16 | ❌ Non specific Cytoplasmic signal in both WT and KO | ❌ Non specific Cytoplasmic signal in both WT and KO |
| **Flow cytometry -** Goat Polyclonal anti-mouse IL33 AF3626 | ✅ IL-33$^+$ fibroblastic stromal cells, lymphatic endothelial cells or oligodendrocytes | No signal in IL-33 KO mice | 1,4,17–19 | ❌ Non specific signal in both WT and KO (~15% positive platelets in both genotypes compared to isotype control) | - |
| **Western blot (WB)** Goat Polyclonal anti-mouse IL33 AF3626 | ✅ 35 kDa band detected in lung lysate/BALF | No 35kDa band in IL-33 KO lung lysate/BALF | 13,14,20–22 | ❌ No 35 kDa band detected | ❌ No 35 kDa band detected |
| **Citrine reporter (IL33-Citrine)** | ✅ IL-33- Citrine$^+$ BM stromal cells | No signal in IL-33-Citrine$^{-/-}$ mice | 2 | ❌ No Citrine$^+$ signal | ❌ No Citrine$^+$ signal |

** The reference numbers correspond to those listed in the Supplemental References section*

1,7–17

1,5,18–20

14,15,21–23

**Supplemental Table 2. Platelet protein modulation in response to IL-33 deficiency or treatment**

| Function Category | Proteins (selected) | IL-33 KO vs WT | IL-33 treated vs WT |
|---|---|---|---|
| **Adhesion – Platelet receptor** | Mmrn1 | ↓ | |
| **Cytoskeleton / Shape change** | Flna, Myh9 | ↓ | ↓ |
| **Signaling / Activation** | Psl1, Ptpn11, Lims1, Plcg2 | ↑ | ↓ |
| **Cytoskeleton/Lamelipodium assembly** | Coro1b,Coro1c,Rac1,Rap1b, Plek, Cdc42, Rhoa | ↑ | – |
| **Inflammation / Immune modulation** | Ppbp, Lgals3, Itgam, Mmp12, Anxa1,Anxa2 | – | ↑ |
| **Coagulation / Fibrinolysis** | Gp1ba, Gp1bb, Gp5, F2 | – | ↓ |

#### Supplemental Table 3. List of antibodies

| Marker | Conjugate | Dilution | Clone | Host/Isotype | Supplier | Catalog No. |
|---|---|---|---|---|---|---|
| Anti-Goat | HRP | 1:10000 | | Donkey Polyclonal | Promega | V8051 |
| Anti-goat IgG | Cy3 | 1:2000 | | Bovine Polyclonal | Jackson ImmunoResearch | 805165180 |
| CD11b | FITC | 1:2000 | M1/70 | Rat IgG2b, κ | eBiosciences | 11-0112 |
| CD11c | FITC | 1:300 | N418 | Armenian Hamster IgG | eBiosciences | 11-0114 |
| CD127 | ef450 | 1:100 | A7R34 | Rat IgG2a, κ | eBiosciences | 48-1271-82 |
| CD150 | PEcy5 | 1:500 | TC15-12F12.2 | Rat IgG2a, λ | Biolegend | 115911 |
| CD16/32 | | 1:1000 | | | | |
| CD19 | FITC | 1:2000 | eBioAd3 | Rat IgG2a, κ | eBiosciences | 11-0193 |
| CD3 | FITC | 1:500 | 17A2 | Rat IgG2b, κ | Biolegend | 100203 |
| CD4 | FITC | 1:1000 | GK1.5 | Rat IgG2b, κ | eBiosciences | 11-0041-85 |
| CD40L (CD154) | PerCP-Cy5.5 | 1:250 | MR1 | Armenian Hamster IgG | Biolegend | 106514 |
| CD41 | BV786 | 1:500 | MWReg30 | Rat IgG1, κ | BD Biosciences | 740903 |
| CD42b (GpIbα) | DL649 | 1:500 | Xia.G5 | Rat IgG2b | Emfret | M040-3 |
| CD45 | AF700 / APC Fire810 | 1:1000 | 30-F11 | Rat IgG2b, κ | Biolegend | 103128 / 103174 |
| CD45R/B220 | FITC | 1:300 | RA36B2 | Rat IgG2a, κ | BD Biosciences | 553088 |
| CD48 | BV711 | 1:500 | HM48-1 | Armenian Hamster IgG | Biolegend | 103439 |
| CD49b | FITC | 1:100 | Sam G4 | Rat IgG2b | Emfret | M070-1 |
| CD49f | FITC | 1:500 | GoH3 | Rat IgG2a, κ | Biolegend | 313605 |
| CD61 | AF647 | 1:500 | 2C9.G2 | Armenian Hamster IgG | Biolegend | 104314 |
| CD63 | APC Cy7 | 1:500 | NVG-2 | Rat IgG2a, κ | Biolegend | 143907 |
| CD9 | BV605 | 1:250 | KMC8 | Rat IgG2a, κ | BD Biosciences | 740360 |
| CD90.2 | APCFire750 | 1:100 | 53-2.1 | Rat IgG2a, κ | Biolegend | 140326 |
| ckit | BV650 | 1:500 | 2B8 | Rat IgG2b, κ | BioLegend | 105853 |
| CLEC2 | PE | 1:250 | 17D9/CLEC-2 | Rat IgG2a, κ | Biolegend | 146104 |
| FCeRI | FITC | 1:100 | MAR-1 | Armenian Hamster IgG | eBiosciences | 11-5898-85 |
| GPVI | FITC | 1:100 | JAQ1 | Rat IgG2a | Emfret | M011-1 |
| Gr1 | FITC | 1:100 | RB6-8C5 | Rat IgG2b, κ | Biolegend | 108406 |
| IL-33 | - | 1:200 | - | Goat polyclonal | R&D Systems | AF3626 |
| NK1.1 | FITC | 1:300 | PK136 | Mouse IgG2a, κ | Biolegend | 108706 |
| P-Selectin | BV711 | 1:500 | RB40.34 | Rat IgG1, λ | BD Biosciences | 740693 |
| Sca1 | APC | 1:500 | D7 | Rat IgG2a, κ | eBiosciences | 17-5981-83 |
| ST2 | PECy7 | 1:200 | RMST2-2 | Rat IgG2a, κ | Invitrogen | 25-9335-82 |
| ST2 Isotype Ctrl | - | 1:200 | IgG2a κ | Rat IgG2a, κ | Invitrogen | 25-4321-82 |
| Ter119 | APC / PECy7/FITC | 1:500 | Ter119 | Rat IgG2b, κ | Biolegend | 116212 / 116221/115921 |
| Viability | RedDye | 1:500 | - | - | Cytek | SKU R7-60008 |

**Supplemental Table 4. List of reagents**

| Reagent | Supplier | Catalog/Ref No. |
|---|---|---|
| ACD | Sigma | C3821 |
| ADP (bacterial) | Sigma | A2754 |
| Apyrase | Sigma | A6535 |
| BSA | Sigma | A2153 |
| $CaCl_2 \cdot 6H_2O$ | Sigma/Merck | |
| $CaCl_2 \cdot 6H_2O$ | Sigma/Merck | 442909 |
| Collagen I | BD | 354236 |
| CRP-XL | CambCol | - |
| D-glucose monohydrate | Sigma | D9559 |
| DiOC6 | ThermoFisher | D273 |
| EDTA 0.5M | Euromedex | EU0084-B |
| Elisa Duoset Dkk1 | RnD | DY1765 |
| Elisa Duoset PF4 | RnD | DY595 |
| Elisa Duoset sCD40L | RnD | DY1163 |
| Fibrinogen | Invitrogen | RP-43142 |
| Fibrinogen (soluble) (AF488) | Invitrogen | F13191 |
| Fibronectin (human) | CORNING | 354008 |
| FOXP3 Fix/Perm Kit | Invitrogen eBioscience | 00-5523-00 |
| Heparin | Sigma | H3149 |
| HEPES | Sigma | H4034 |
| KCl | Sigma/Merck | P3911 |
| Lab Tek - Chamber slides | ThermoFisher | 178599 |
| Laminin | Sigma/Merck | L4544 |
| $MgCl_2 \cdot 6H_2O$ | Sigma/Merck | M2393 |
| Microcapillaries flow chamber | Cellix | Vena8 Fluoro+ |
| Minivette 50 µL | SARSTEDT | 17.2113.050 |
| NaCl | Sigma | 59625 |
| $NaH_2PO_4$ | Sigma | S3139 |
| $NaHCO_3$ | Sigma | S7277 |
| $PGI_2$ | Abcam | ab120912 |
| Phalloidin-CF488A | Biotium | 00042-T |
| Pipes | Sigma | P6757 |
| Podoplanin (Fc chimera) | Biolegend | 551704 |
| Precision Count Beads | Biolegend | 424902 |
| Thrombin | Sigma-Aldrich/Merck | T6884 |
| Tiazole Orange | Sigma | 390062 |
| Vitronectin (human) | Sigma/Merck | SRP3186 |

**Supplemental Table 5. List of upregulated protein in platelets from IL33 KO mice**

**Supplemental Table 6. List of downregulated protein in platelets from IL33 KO mice**

**Supplemental Table 7. List of upregulated protein in platelets from IL33 treated mice**

**Supplemental Table 8. List of downregulated protein in platelets from IL33 treated mice**

**Supplemental Figure 1**

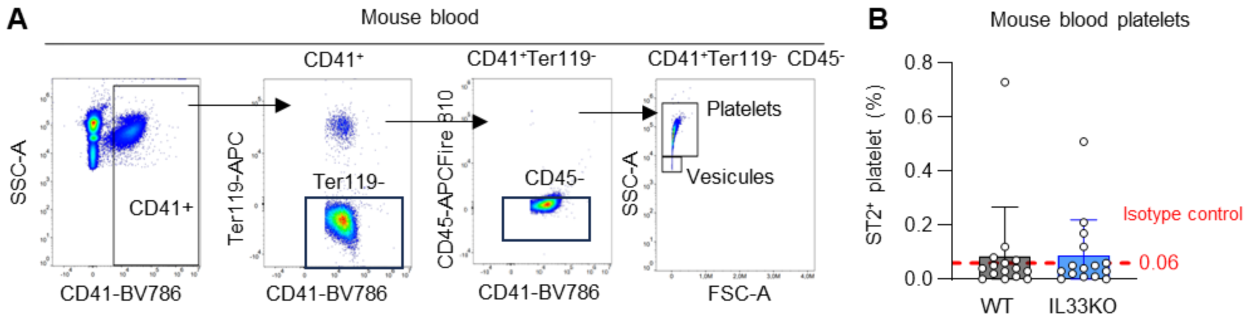


**Supplemental Figure 1. ST2 is not expressed by blood platelets** (A) Representative blood platelet gating strategy. (B) Quantification of the percentage of platelets expressing ST2 in WT and IL-33-deficient mice (n = 14 per group) using isotype control as negative reference.

### Supplemental Figure 2

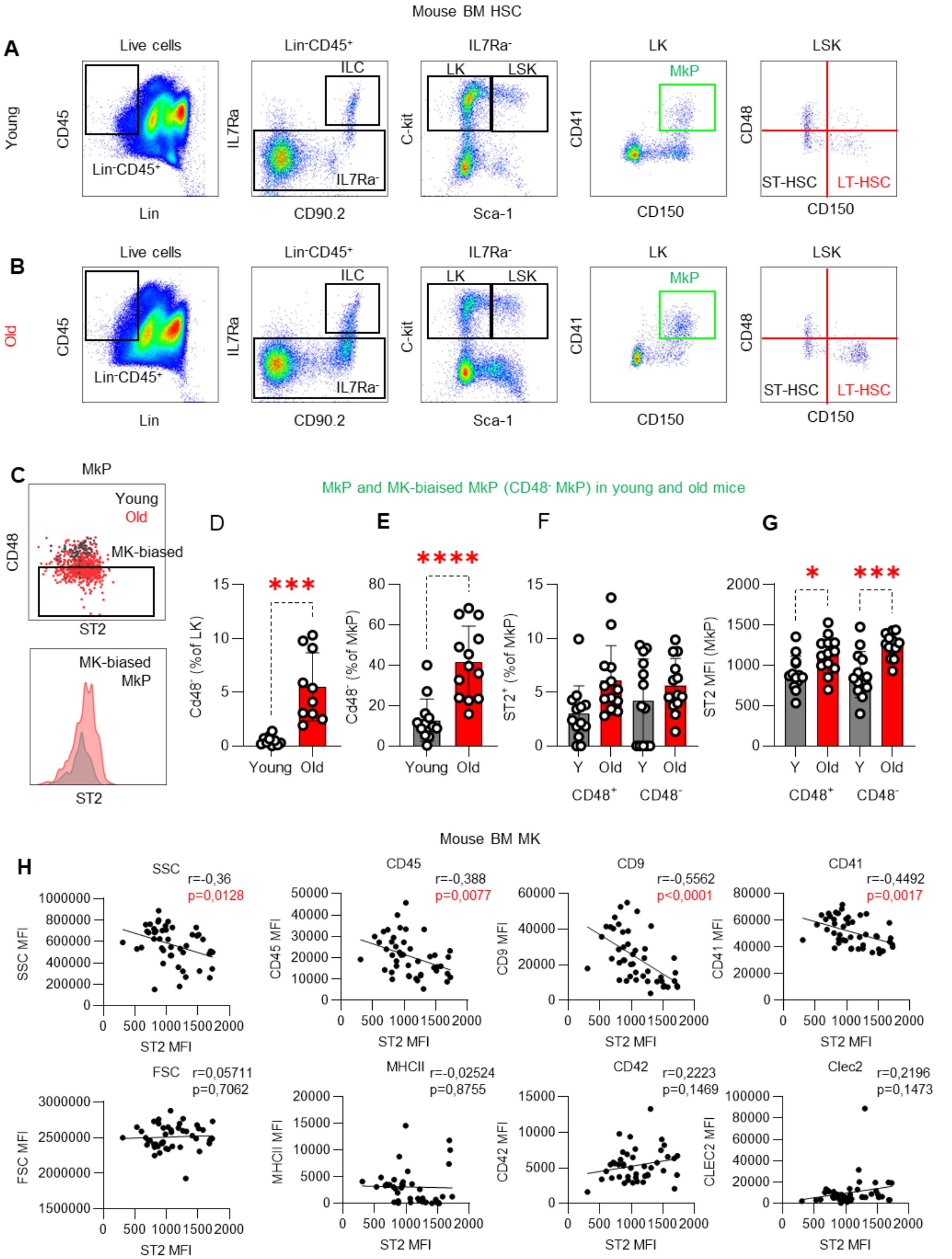


**Supplemental Figure 2. Gating strategies and ST2 expression analysis in hematopoietic and megakaryocyte progenitors from mouse bone marrow.**

(A-B) Gating strategy for identifying long-term hematopoietic stem cells (LT-HSCs) and megakaryocyte progenitors (MkPs) in the bone marrow of young (8-week-old) and aged (2-year-old) mice. (C) Gating strategy for identifying megakaryocyte-biased MkPs (CD48$^{-}$ MkPs) and ST2 expression on this population. (C-D) Quantification of MK-biased MkPs in young and aged mice. (E-G) Quantification of ST2$^{+}$ cells and ST2 expression levels within MkPs. Data obtained from n = 13 mice per group (biological replicates) across three independent experiments conducted on separate collection and analysis days. Investigators were blinded to mouse age during bone marrow collection and analysis. (H) Correlation between ST2 mean fluorescence intensity (MFI) and megakaryocyte (MK) markers in bone marrow MKs (Lin$^{-}$CD45$^{-}$CD41$^{+}$FSC$^{Hi}$) from young and old mice. The Pearson correlation coefficient (r) and two-tailed p-value are indicated. Data represent n=46 mice collected across 5 different experiments.

**Supplemental Figure 3**

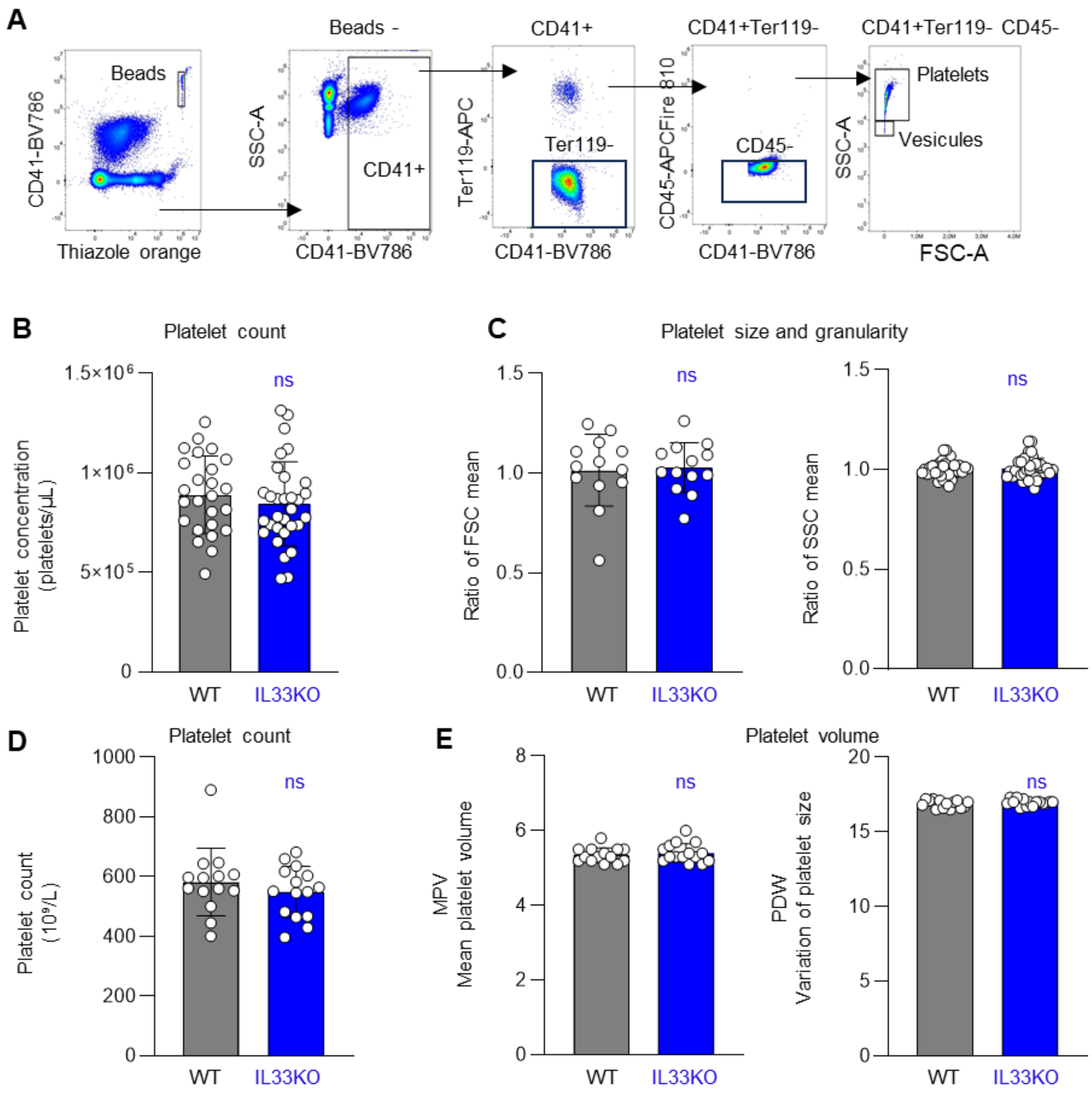


**Supplemental Figure 3. Total IL-33 deficiency does not affect platelet count or morphology**

(A) Gating strategy used to identify platelets from diluted whole blood by flow cytometry. (B–C) Quantification of platelet count (B) and analysis of platelet morphology (size and granularity) (C) by flow cytometry in whole blood from WT and IL33KO mice. Data are pooled from 4 independent experiments; each point represents one mouse. (D–E) Platelet count (D) and morphological parameters (mean platelet volume and platelet size distribution) (E) assessed using a hematology analyzer in WT and IL33KO mice. Data represent n= 25-31 mice per group collected from 3 independent experiments. Statistical significance was determined using a t-test.

#### Supplemental Figure 4

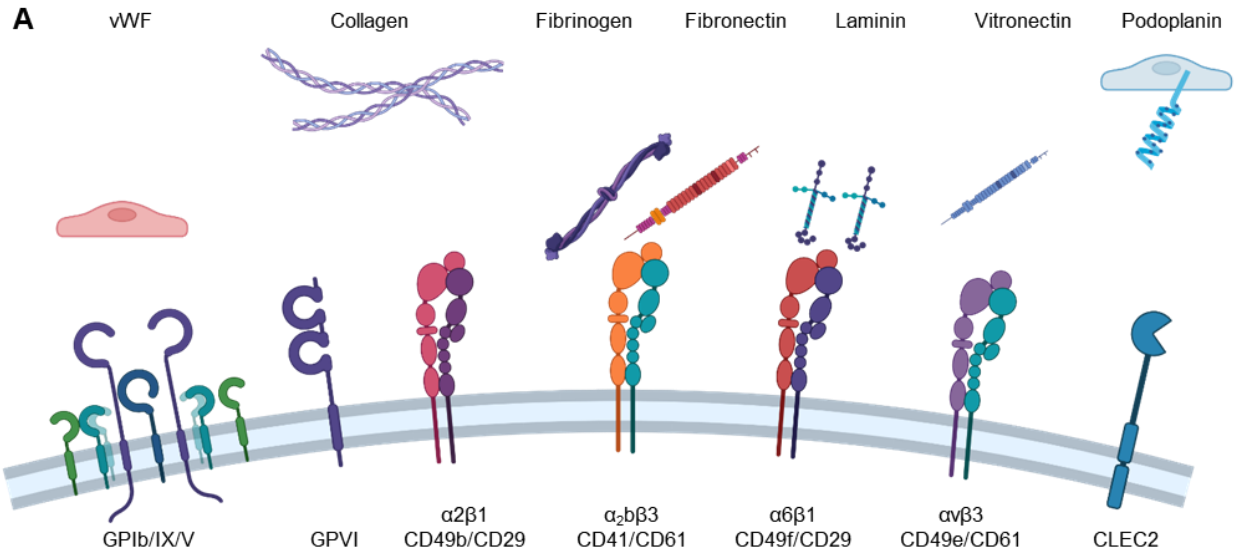


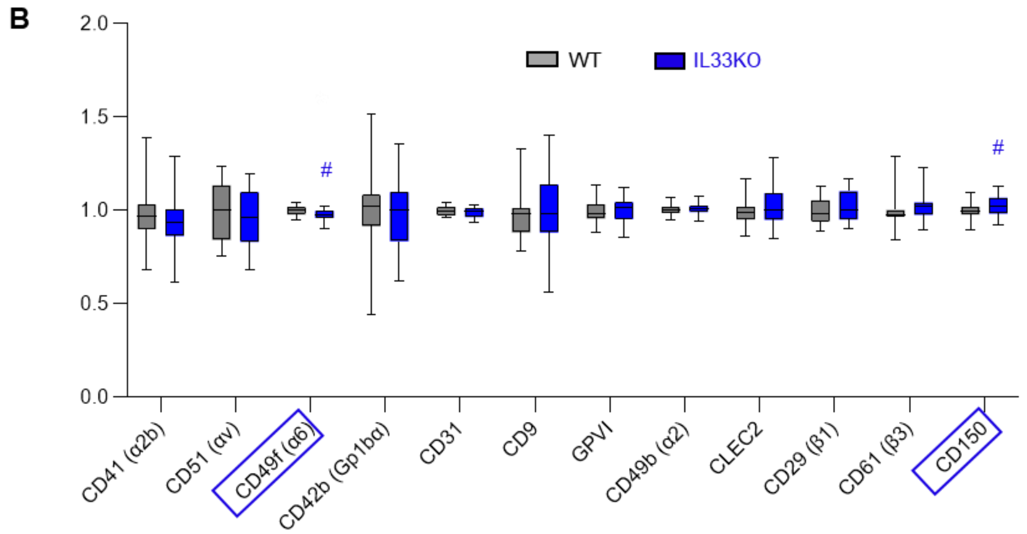


**Supplemental Figure 4. Expression of integrins and surface receptors in platelets from IL-33-deficient (IL33KO) and wild-type (WT) mice**

(A) Schematic representation of platelet integrins and receptors involved in interactions with non-soluble activators, including components of the extracellular matrix, activated endothelium, and lymphatic vessels. (B) Quantification of integrin and receptor expression levels on platelets from WT and IL33KO mice. Data represent the distribution (box-and-whisker plots) of n = 28–31 mice pooled from five independent experiments. Statistical analysis was performed using a t-test.

#### Supplemental Figure 5

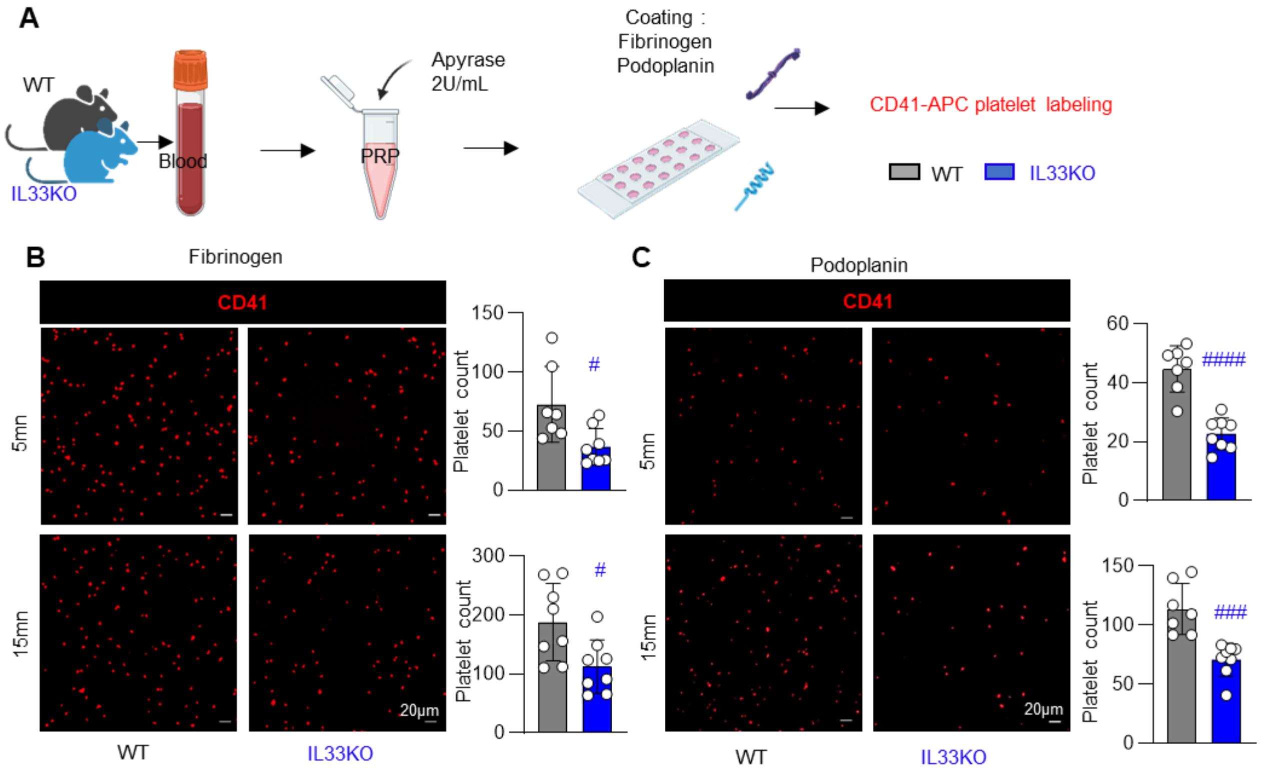


**Supplemental Figure 5. Platelets from IL33KO mice have a fixation defect on fibrinogen and podoplanin with apyrase inhibition during the deposit**

(A) Schema of fixation assays analysed by immunofluorescence of platelets from WT or IL33KO mice. (B) Representative images and quantitative analysis of adherent and spreading PRP from WT or IL33KO mice at 5, 15 mn with Apyrase (2U/mL) inhibition during the deposit on fibrinogen (B) and on podoplanin (C). Platelets are labelled with CD41-APC. Quantitative data represent n = 7–8 mice per group from two pooled independent experiments. t-test, #<0,05; ##<0,01; ###<0,001;####<0,0001

#### Supplemental Figure 6

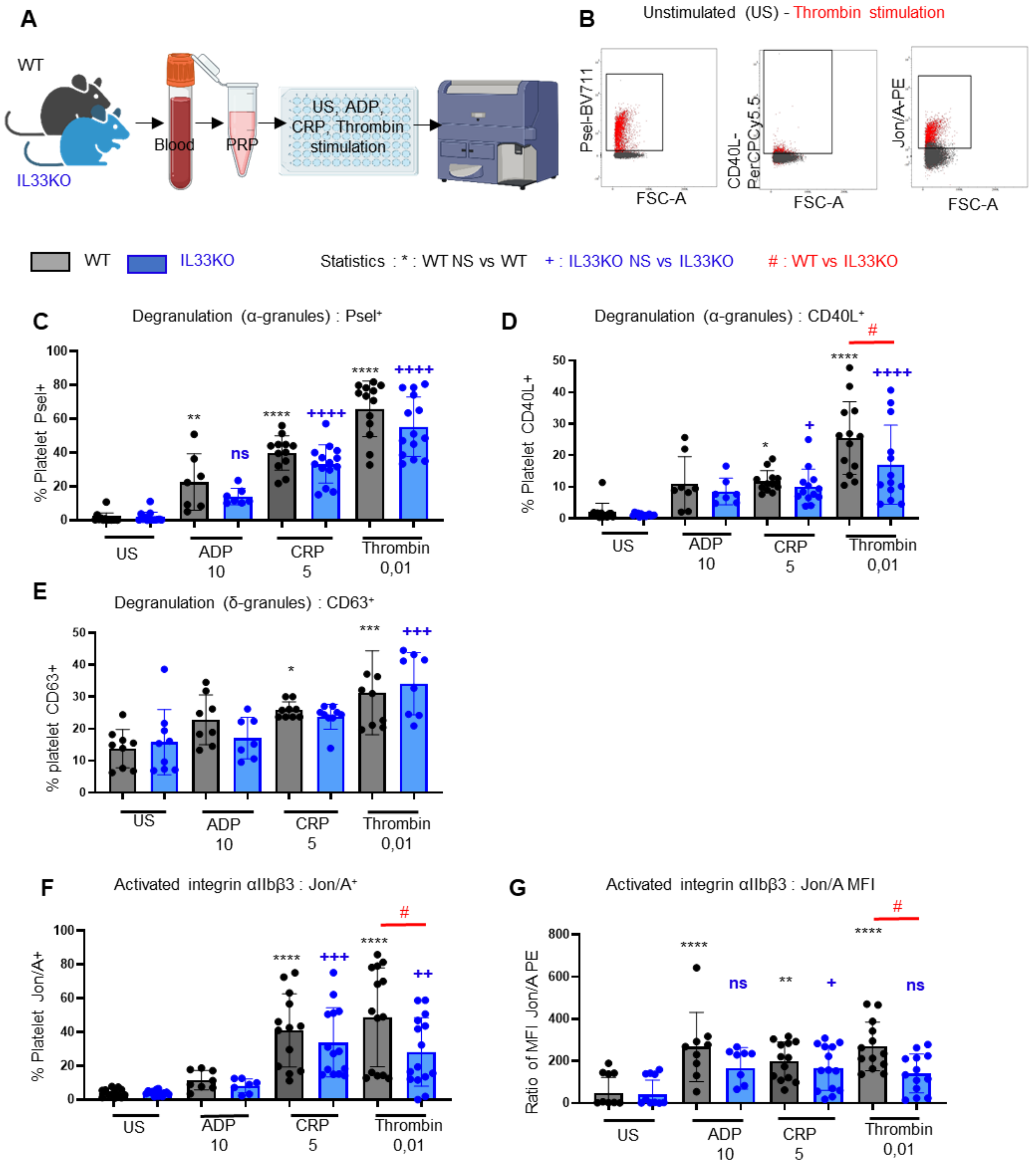


**Supplemental Figure 6. IL33KO platelets present mild defects in degranulation and integrin activation**

(A) Schematic and (B) gating strategy for analyzing platelet activation markers by flow cytometry in PRP from WT or IL33KO mice, either unstimulated (US) or after 15 minutes of *in vitro* stimulation with ADP (10µM), CRP (5µg/mL), and thrombin (0.01 U/mL). Analysis of the percentage of platelets showing surface expression of P-selectin (C), CD40L (D), CD63 (E), Jon/A (F), and Jon/A fluorescence intensity levels (G). Data represent n=7-14 mice per group from three pooled independent experiments. Statistical significance was determined by one-way ANOVA with Sidak's multiple comparisons test, $*p < 0.05$; $**p < 0.01$; $***p < 0.001$; $****p < 0.0001$.

#### Supplemental Figure 7

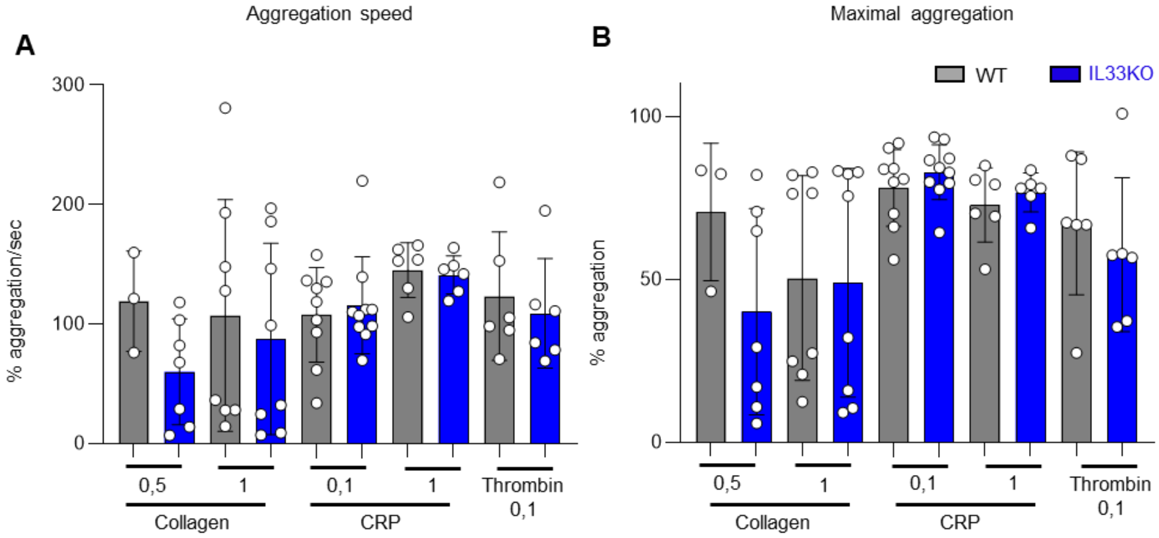


**Supplemental Figure 7. Platelets from IL33KO mice exhibit normal aggregation responses to collagen, CRP, and thrombin stimulation compared to WT platelets**

(A) Aggregation kinetics of platelets from WT and IL33KO platelet-rich plasma (PRP) following stimulation with collagen (0.5 or 1 µg/mL), CRP (0.1 or 1 µg/mL), or thrombin (0.1 IU/mL). (B) Maximal platelet aggregation capacity in WT and IL33KO PRP under the same stimulation conditions. n= 3-10 mice collected from 2 independent experiments. Statistical analysis was performed using one-way ANOVA followed by Sidak's multiple comparisons test.

### Supplemental Figure 8

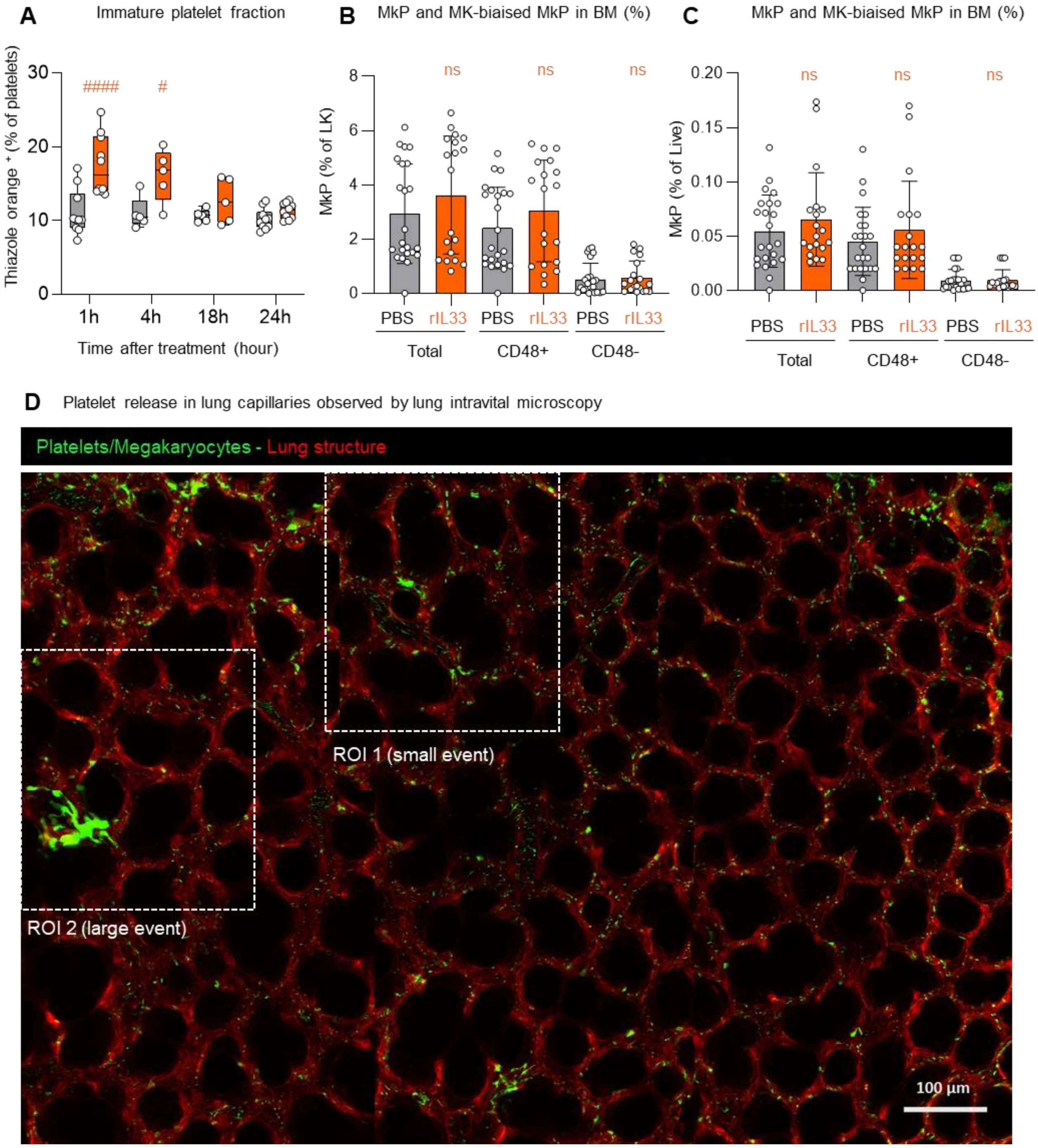


**Supplemental Figure 8. Platelet release within pulmonary capillaries visualized by intravital lung microscopy**

(A) Quantification of the immature platelet fraction (percentage of thiazole orange positive platelets) in blood collected 1 h, 4 h, 18 h, or 24 h after intranasal rIL-33 treatment (1 µg). n = 4–10 mice from 2 independent experiments. (B–C) Quantification of bone marrow megakaryocyte progenitor (MkP) populations, including canonical ($CD48^{+}$) and non-canonical ($CD48^{-}$) subsets, following treatment with recombinant IL-33 (rIL-33; 3 × 1 µg, intranasal). Data are expressed as (B) a percentage of the parent population ($LK=CD45^+Lin^-ckit^+Sca1^-$) or (C) as a percentage of live cells. Data obtained from n = 13 mice per group (biological replicates) across three independent experiments conducted on separate collection and analysis days. Investigators were blinded to mouse age during bone marrow collection and analysis. (D) Representative image from intravital microscopy of the PF4-mTmG lung showing $PF4^+$ platelets in the pulmonary capillaries, small proplatelets (ROI1) and larger proplatelet/megakaryocyte structures (ROI2) actively releasing platelets within pulmonary capillaries. Time lapse images of regions of interest (ROI1 and ROI2) are shown at higher magnification in Figure 6H. Scale bar = 100 µm.

**Supplemental Figure 9**

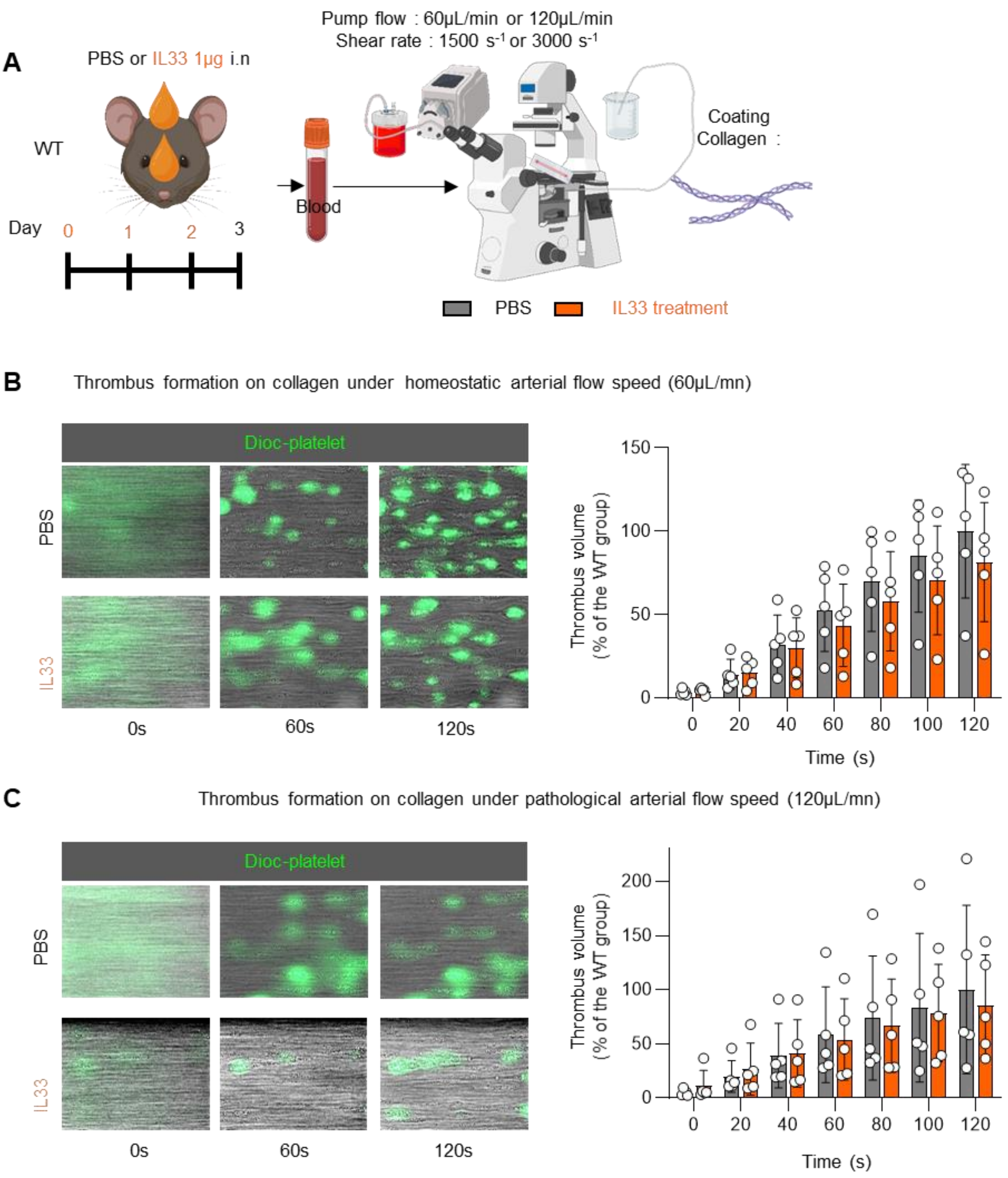


**Supplemental Figure 9. IL-33 treatment does not alter thrombus volume under arterial flow conditions**

(A) Schematic representation of the thrombus formation assay on collagen under arterial flow using whole blood from WT mice treated with PBS or recombinant human IL-33 (rIL-33h). (B) Representative images and quantification of thrombus volume (expressed as percentage relative to the mean value of the PBS-treated WT group) under homeostatic arterial flow conditions (60 µL/min). (C) Representative images and quantification of thrombus volume under pathological arterial flow conditions (120 µL/min), using the same analysis method. Data are representative of 3 independent experiments (n = 5 mice per group). Statistical analysis was performed using two-way ANOVA followed by Sidak's multiple comparisons test; #$p < 0.05$. Investigators were blinded to treatment during the thrombus formation assay.

**Supplemental Figure 10**

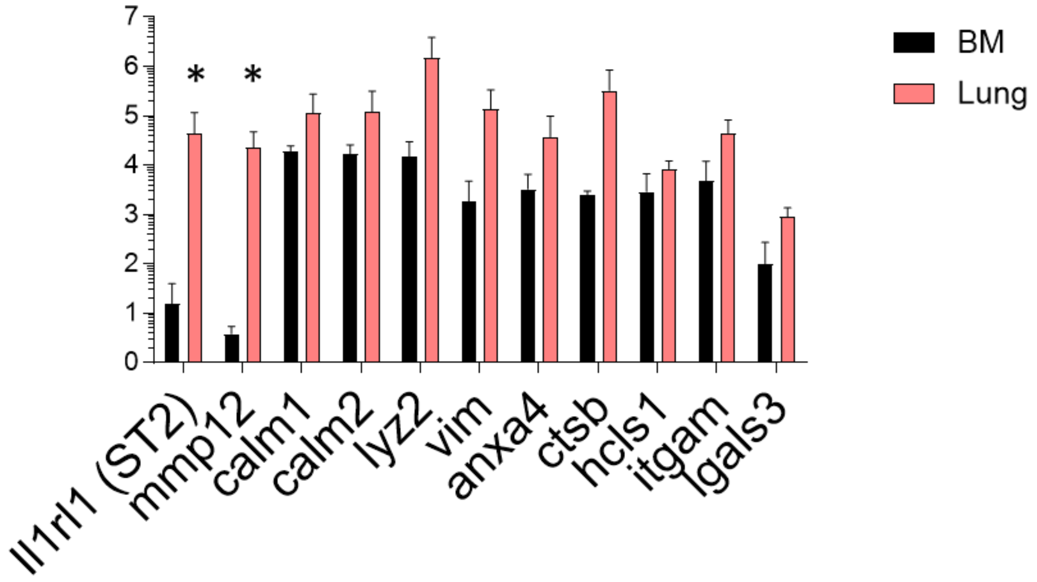


**Supplemental Figure 10. Gene expression in lung versus bone marrow megakaryocytes for ST2 and proteins upregulated in platelets from mice treated with rIL-33 *in vivo*.**

Comparative RNA-seq analysis of murine bone marrow and lung megakaryocytes (PF4-nTnG/CD41$^+$ cells). ST2 and abundant proteins upregulated in platelets from mice treated with rIL-33 *in vivo* are shown. Data from Lefrançais *et al*., Nature, 2017 (n=3 mice per group)

#### Supplemental Video legend

##### Supplemental Video 1. Rapid platelet release within pulmonary capillaries visualized by intravital lung microscopy

60-minute time-lapse intravital microscopy of the PF4-mTmG mouse lung showing $PF4^{+}$ platelets circulating in pulmonary capillaries, along with small proplatelets and larger proplatelet/megakaryocyte structures actively releasing platelets. This process, corresponding to Figure 6H and Supplemental Figure 7, highlights the rapid fragmentation of large megakaryocytes into individual platelets within 30–40 minutes, significantly faster than in the bone marrow.
